# Microbial community shifts associated with the ongoing stony coral tissue loss disease outbreak on the Florida Reef Tract

**DOI:** 10.1101/626408

**Authors:** Julie L. Meyer, Jessy Castellanos-Gell, Greta S. Aeby, Claudia Häse, Blake Ushijima, Valerie J. Paul

## Abstract

As many as 22 of the 45 coral species on the Florida Reef Tract are currently affected by stony coral tissue loss disease (SCTLD). The ongoing disease outbreak was first observed in 2014 in Southeast Florida near Miami and as of early 2019 has been documented from the northernmost reaches of the reef tract in Martin County down to Key West. We examined the microbiota associated with disease lesions and apparently healthy tissue on diseased colonies of *Montastraea cavernosa, Orbicella faveolata, Diploria labyrinthiformis*, and *Dichocoenia stokesii*. Analysis of differentially abundant taxa between disease lesions and apparently healthy tissue identified five unique amplicon sequence variants enriched in the diseased tissue in three of the coral species, namely an unclassified genus of Flavobacteriales and sequences identified as *Fusibacter* (Clostridiales), *Planktotalea* (Rhodobacterales), *Algicola* (Alteromonadales), and *Vibrio* (Vibrionales). In addition, several groups of likely opportunistic or saprophytic colonizers such as Epsilonbacteraeota, Patescibacteria, Clostridiales, Bacteroidetes, and Rhodobacterales were also enriched in SCTLD disease lesions. This work represents the first microbiological characterization of SCTLD, as an initial step toward identifying the potential pathogen(s) responsible for SCTLD.

## INTRODUCTION

An ongoing outbreak of coral disease termed stony coral tissue loss disease (SCTLD) is currently impacting at least twenty coral species on the Florida Reef Tract (Florida Keys National Marine Sanctuary, 2018; Lunz et al., 2017). Outbreak levels of disease were first observed in late 2014 off Virginia Key, FL, and over the next year the disease was detected both north and south along the Florida Reef Tract (Precht et al., 2016). Active monitoring by county, state, and federal agencies in Southeast Florida and the Florida Keys National Marine Sanctuary has since recorded disease occurrence from northern Martin County south and west through the Florida Reef Tract to Key West. The geographic extent, the number of coral species impacted, and the modeling of disease incidence all support the conclusion that this is a highly contagious disease (Muller et al., 2018).

Monitoring efforts and early studies indicate that not all coral species are equally susceptible to SCTLD, with corals like *Dendrogyra cylindrus, Dichocoenia stokesii*, and *Meandrina meandrites* succumbing very rapidly to the disease (Florida Keys National Marine Sanctuary, 2018). Important reef-building species like *Montastraea cavernosa* and *Siderastrea siderea* have also been heavily impacted (Walton et al., 2018). In addition to variability in response among different coral species, environmental factors may influence disease severity. For example, the highest prevalence of SCTLD in Southeast Florida during surveys from 2012 to 2016 also coincided with the highest prevalence of coral bleaching (Walton et al., 2018) in response to elevated sea surface temperatures in the summer and winter of 2014 (Manzello, 2015). In addition, dredging operations between 2013 and 2015 in the channel at the Port of Miami were correlated with increased sedimentation and partial coral mortality near the site where the disease outbreak was detected (Miller et al., 2016). Responses within species may also be constrained by local reef conditions. In the Upper Keys, monitored colonies of *S. siderea* and *Pseudodiploria strigosa* experienced higher mortality from white plague-like disease (prior to the onset of SCTLD incidence) in offshore reefs dominated by octocorals and macroalgae in comparison to inshore patch reefs with higher coral cover (Rippe et al., 2019).

Currently, it is unclear whether all of the impacted species are suffering from the same disease, and there are no existing diagnostic tools to positively identify SCTLD, as is the case for most coral diseases. Several enigmatic tissue loss diseases, also known as white syndromes as they are characterized by the white skeleton left exposed by the disease, have been identified in corals around the world, most without the identification of a definitive causative agent (Mera and Bourne, 2018). In the Caribbean, potential pathogens have been identified for three coral tissue loss diseases: white band disease type II, white plague disease type II, and white pox disease. White band disease type II has been attributed to *Vibrio harveyi* (Gil-Agudelo et al., 2006; *Ritchie and Smith, 1998) and is part of the suite of white band diseases that contributed to the drastic decline in Acropora palmata* and *A. cervicornis* in the 1980s (Aronson and Precht, 2001). White plague disease type II has also reportedly impacted dozens of other Caribbean stony coral species (Weil et al., 2006), and an outbreak of this disease in the 1990s was attributed to an alphaproteobacterium (Richardson et al., 1998), and the potential pathogen, named *Aurantimonas coralicida*, was subsequently isolated from *Dichocoenia stokesii* (Denner et al., *2003*). *A later study using both traditional clone libraries of small subunit ribosomal genes and microarray assays of Caribbean Orbicella faveolata* affected by white plague disease type II failed to detect the putative pathogen *Aurantimonas coralicida* (Sunagawa et al., 2009). *White pox disease (acroporid serratiosis), which exclusively impacts Acropora palmata* corals in the Caribbean, has been attributed to the opportunistic bacterial pathogen *Serratia marcescens* (Patterson et al., 2002), which may have originated from human sewage (Sutherland et al., 2010). In contrast, putative pathogens have not been identified for most common coral diseases (Weil et al., 2006).

To aid in the discovery of the pathogen or pathogens responsible for the current SCTLD outbreak, we characterized the microbiomes associated with four coral species with SCTLD from sites in Southeast Florida and the Middle Keys. For each diseased colony, we examined microbiome composition at the disease lesion, in apparently healthy polyps immediately adjacent to the lesion, and in apparently healthy polyps far from the disease lesion. Neighboring corals that had no apparent disease lesions were also sampled when available. This strategy was used to determine the microbiome shifts associated with SCTLD, which may contribute to a better understanding of the disease ecology of the current devastating outbreak on the Florida Reef Tract.

## MATERIALS AND METHODS

Coral colonies with SCTLD were identified in Southeast Florida offshore from Ft. Lauderdale and in the Middle Florida Keys near Long Key Bridge. *Montastraea cavernosa* and *Orbicella faveolata* were sampled from SE Florida in July and December 2017. *Diploria labyrinthiformis* and *Dichocoenia stokesii* were sampled in the Middle Keys in December 2017. For *D. labyrinthiformis*, and *D. stokesii*, the disease lesions were peripheral or apical, with multiple lesions often proceeding to coalescence (Figure 1). In all sampled *M. cavernosa* and *O. faveolata*, lesions were clearly preceded by a row of bleached and/or partially bleached polyps (Figure 1A).

**Figure 1.**
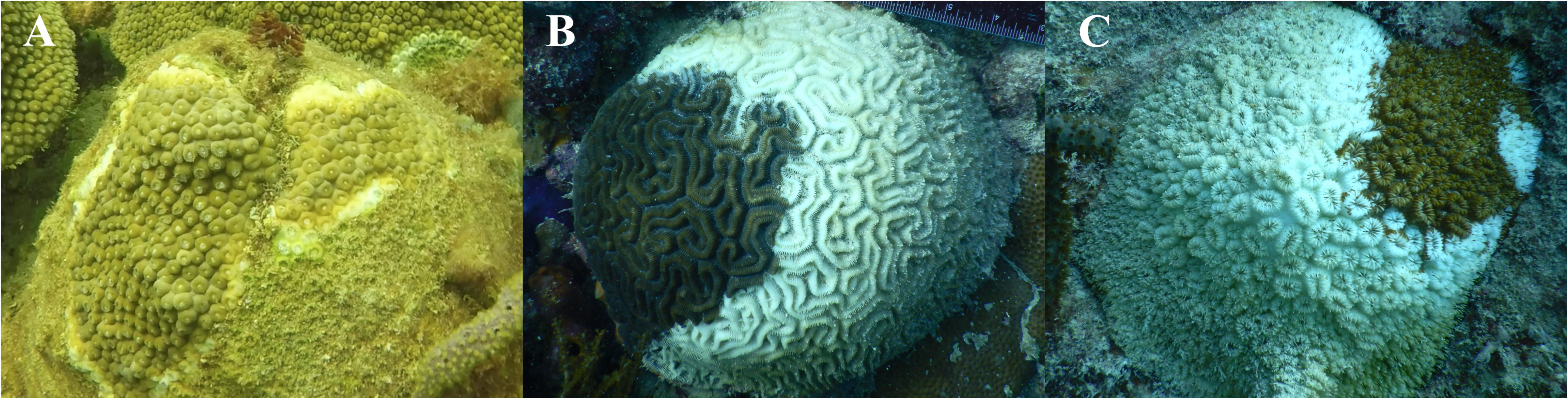
Stony coral tissue loss disease appearance in three coral species from the Florida Reef Tract: *Montastraea cavernosa* (A), *Diploria labyrinthiformis* (B), and *Dichocoenia stokesii* (C).

Coral mucus/tissue samples were collected while SCUBA diving using agitation of the coral surface and aspiration with a needleless syringe. Care was taken to collect all of the released particulates via the syringe to prevent the spread of diseased material. Samples were collected at the lesion, at the first row of healthy-looking polyps adjacent to the lesion, and as far as possible away from the lesion in apparently healthy tissue from nine colonies of *M. cavernosa* and four colonies each of *D. labyrinthiformis*, and *D. stokesii* (Table S1). Six healthy neighboring corals of *M. cavernosa* and one healthy neighboring coral of *D. labyrinthiformis* were also sampled (Table S1). Samples were kept on ice until reaching shore where mucus and tissue were separated by settling out of the seawater and frozen until extraction of DNA.

Extractions of genomic DNA were performed with a DNeasy Powersoil kit (Qiagen, Germantown, MD) according to the manufacturer’s instructions, with bead beating for 10 min. The V4 region of the 16S rRNA gene was amplified in triplicate for each sample using the 515F (Parada et al., 2016) and 806RB (Apprill et al., 2015) Earth Microbiome primers and thermocycler protocol (Caporaso et al., 2012) in 25-μl reactions containing Phusion High-fidelity Master Mix (New England Biolabs, Ipswich, MA), 0.25μM of each primer, 3% dimethyl sulfoxide (as recommended by the manufacturer of the polymerase), and 2 μl of DNA template. Triplicate samples were consolidated and cleaned with a MinElute PCR purification kit (Qiagen) and quantified with a Nanodrop 1000 (Thermo-Fisher Scientific, Waltham, MA). DNA extraction kit blanks were produced alongside regular DNA extractions without the addition of any starting coral biomass. Two extraction kit blanks and one PCR blank were processed alongside coral samples through sequencing with unique Earth Microbiome barcodes. One final amplicon pool containing 240 ng of each sample library was submitted to the University of Florida Interdisciplinary Center for Biotechnology Research for sequencing. The amplicon pool was quantified using quantitative PCR and QUBIT for quality control and size selected with ELF to produce fragment sizes in the desired range for amplicon sequencing. Sequencing was performed on an Illumina MiSeq with the 2×150bp v. 2 cycle format.

Adapters and primers were removed from raw sequencing reads with cutadapt v. 1.8.1 (Martin, 2011). Sequencing reads are available in NCBI’s Sequence Read Archive under Bioproject PRJNA521988. Further processing of amplicon libraries was completed in R v. 3.5.1. All R scripts and data needed to recreate the figures in this manuscript are available on GitHub (https://github.com/meyermicrobiolab/Stony-Coral-Tissue-Loss-Disease-SCTLD-Project). Quality filtering, error estimation, merging of reads, dereplication, removal of chimeras, and selection of amplicon sequence variants (ASVs) (Callahan et al., 2017) were performed with DADA2 v. 1.6.0 (Callahan et al., 2016), using the filtering parameters truncLen=c(150,150), maxN=0, maxEE=c(2,2), truncQ=2, rm.phix=TRUE to remove all sequences with ambiguous basecalls and phiX contamination. Taxonomy was assigned in DADA2 to ASVs using the SILVA small subunit ribosomal RNA database v. 132 (Yilmaz et al., 2014). The ASV and taxonomy tables, along with associated sample metadata were imported into phyloseq v. 1.22.3 (McMurdie and Holmes, 2013) for community analysis. Sequences that could not be assigned as bacteria or archaea and sequences identified as chloroplasts or mitochondria were removed from further analysis.

ASVs with a mean read count across all samples of less than 5 were removed from the analysis. Zero counts were transformed using the count zero multiplicative method with the zCompositions package in R (Palarea-Albaladejo and Martín-Fernández, 2015). The zero-replaced read counts were transformed with the centered log-ratio transformation, and the Aitchison distance metric was calculated with CoDaSeq (Gloor et al., 2017). Principal component analysis of the Aitchison distance was performed with prcomp in R and plotted with ggplot2 (Wickham, 2016). Analysis of similarity (ANOSIM) and permutational multivariate analysis (PERMANOVA) were performed on the Aitchison distance with vegan (Dixon, 2003) with 999 permutations. Multivariate dispersions of the Aitchison distance were calculated with the betadisper function in vegan and fitted to a linear model to test the significance of coral species and sample condition.

Differential abundance of ASVs between diseased tissue and apparently healthy tissue on diseased colonies was determined with DESeq2 v. 1.20.0 (Love et al., 2014). Differential abundance was assessed within each of the coral species *M. cavernosa, D. labyrinthiformis*, and *D. stokesii*, using the lesion samples and the samples farthest from the lesions. Original count data was used after filtering rows with fewer than 5 counts over the entire row and using the parametric Wald test in DESeq2.

## RESULTS

In the colonies sampled for this study, the disease appeared to progress more slowly in *M. cavernosa* as the exposed skeleton was already colonized by turf algae at the time of collection. In contrast, *D. labyrinthiformis* and *D. stokesii* displayed bare skeleton over most of the colony at the time of sampling, suggesting a more rapid progression of the tissue loss on these highly susceptible species (Figure 1B and 1C). These observations are consistent with reports from the Coral Reef Evaluation and Monitoring Project in Southeast Florida and the Florida Keys (Brinkhuis and Huebner, 2016; Walton et al., 2018).

Microbiomes were characterized from a total of 62 coral samples (Table S1). After quality-filtering and joining, an average of 42,560 reads (2,358 −233,271) per coral sample were used in the analysis. A total of 128 archaeal ASVs and 11,189 bacterial ASVs were detected in coral samples. The three control samples, two from the extraction kit through sequencing and one from PCR through sequencing, were also sequenced and after quality-filtering and joining had an average of 10,144 reads (258-18,348) per control sample, which were classified as 94 bacterial ASVs (Table S2). Overall, microbial community structure differed by coral species (ANOSIM R = 0.407, p = 0.001) (Figure 2). Alpha-and Gammaproteobacteria, Bacteroidetes (Bacteroidia), and Cyanobacteria (Oxyphotobacteria) were commonly detected in all samples (regardless of disease state) from the four coral species (Figure 3), consistent with previous studies of coral microbiomes (Huggett and Apprill, 2018). Amplicon sequence variants (ASVs) classified as Proteobacteria that appeared to be coral mitochondrial sequences based on BLASTn searches were only in *D. stokesii* (Figure 3D). ASVs classified simply as Bacteria in both *D. labyrinthiformis* (Figure 3C), and *D. stokesii* (Figure 3D) were unique sequences, as described in further detail below. Campylobacteria were detected in higher relative abundances in and near disease lesions in *M. cavernosa, D. labyrinthiformis*, and *D. stokesii*, while Deltaproteobacteria were also detected in higher relative abundances in and near lesions of *M. cavernosa* (Figure 3).

**Figure 2.**
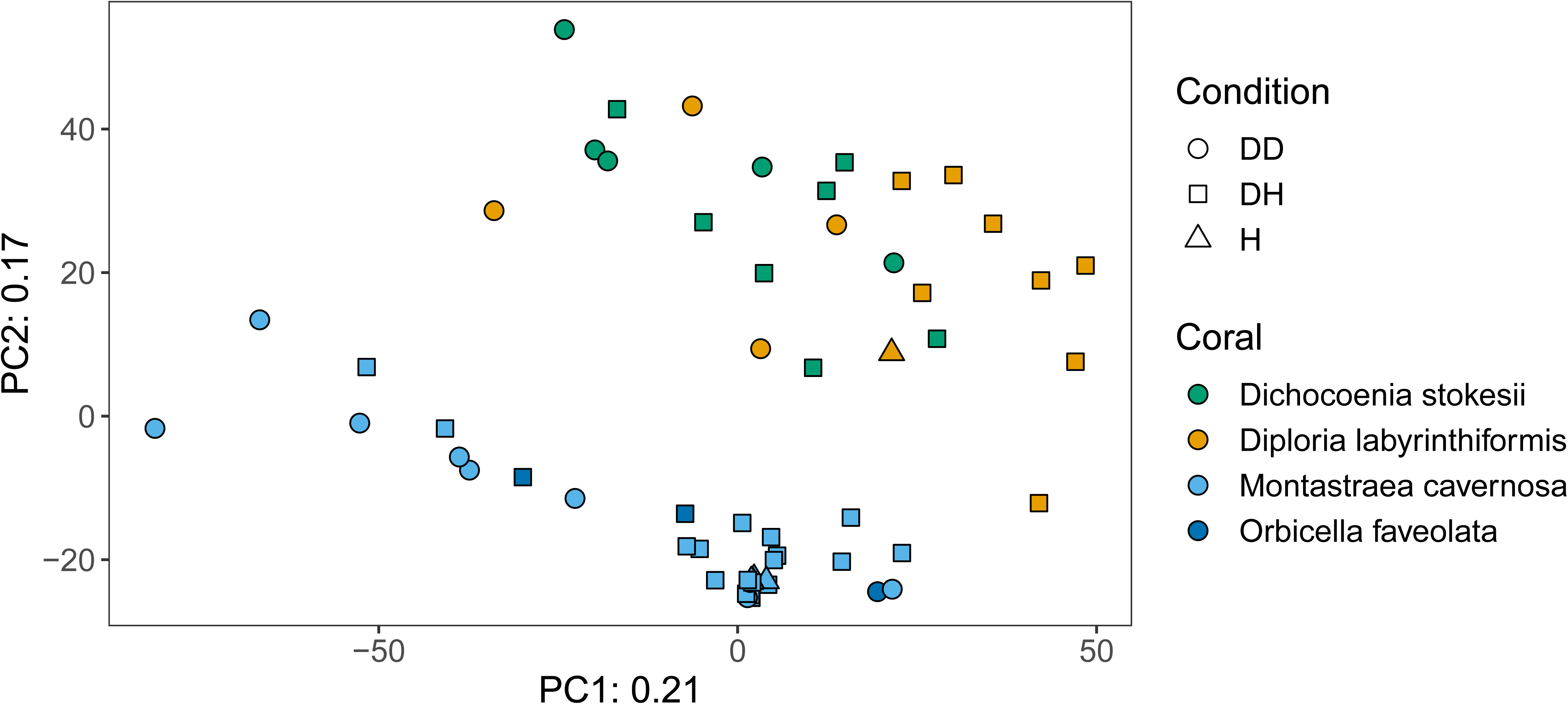
Principal component analysis of microbial community structure in disease lesions (DD), apparently healthy tissue on diseased corals (DH), and apparently healthy neighboring corals with no signs of disease (H).

**Figure 3.**
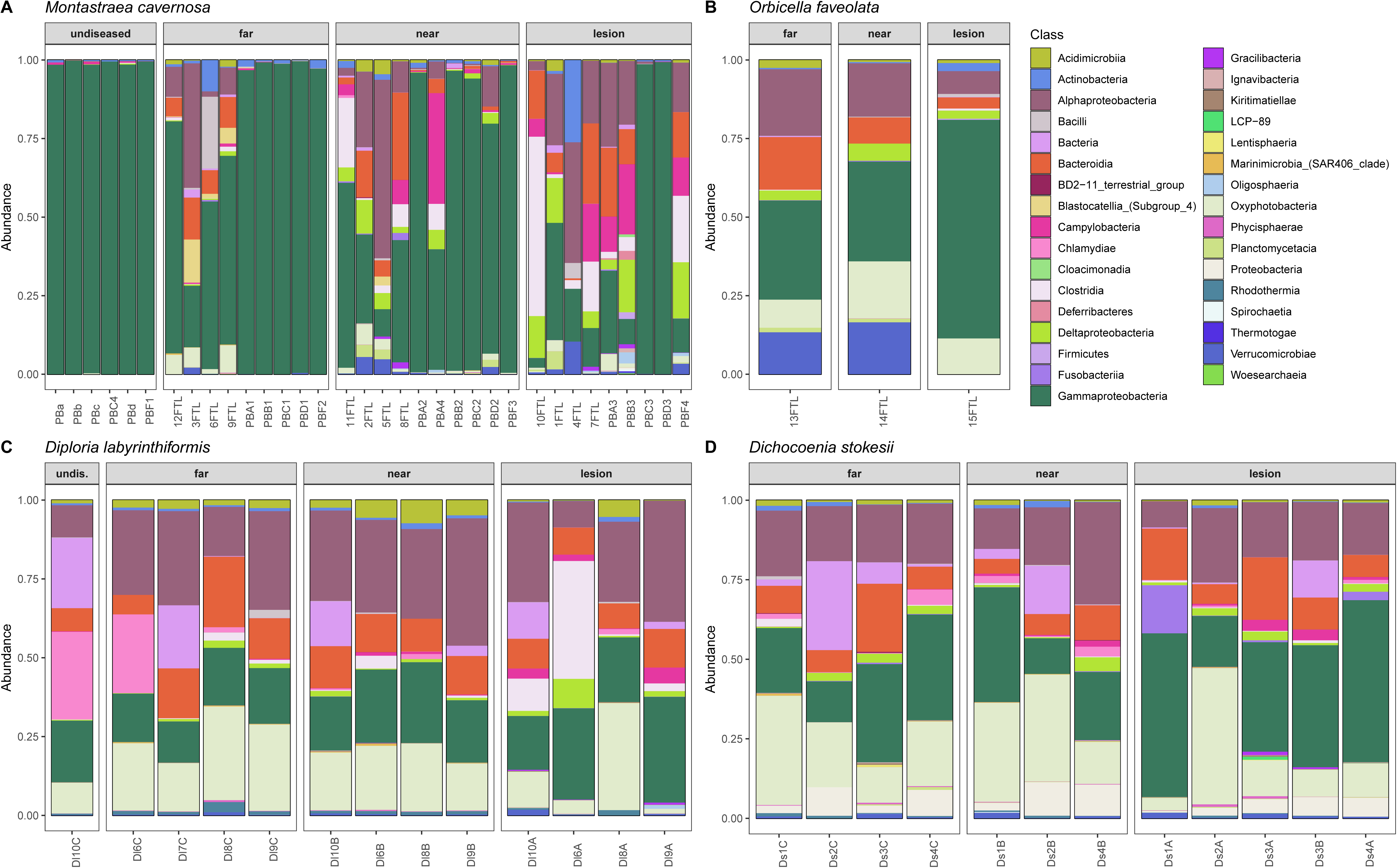
Relative abundance of amplicon sequence variants, colored by Class, in undiseased neighboring corals, apparently healthy tissue far from the disease lesion, apparently healthy tissue near the disease lesion, and disease lesions in *Montastraea cavernosa* (A), *Orbicella faveolata* (B), *Diploria labyrinthiformis* (C), and *Dichocoenia stokesii* (D) with stony coral tissue loss disease.

Disease often has a stochastic effect on microbiome composition (Zaneveld et al., 2017), therefore the dispersion of beta diversity was examined according to health condition (Figure 4). Neighboring healthy colonies were sampled as available and only *M. cavernosa* had a sufficient number of sampled healthy colonies for comparison. In *M. cavernosa*, there is a pattern of health state and stochastic effects on microbiome composition, as healthy colonies had lower variation in their microbiomes, while diseased tissue and apparently healthy tissue on diseased colonies had higher variability in their microbiome composition. In contrast, beta diversity between disease lesions and apparently healthy tissue on diseased colonies of all four coral species were similarly variable.

**Figure 4.**
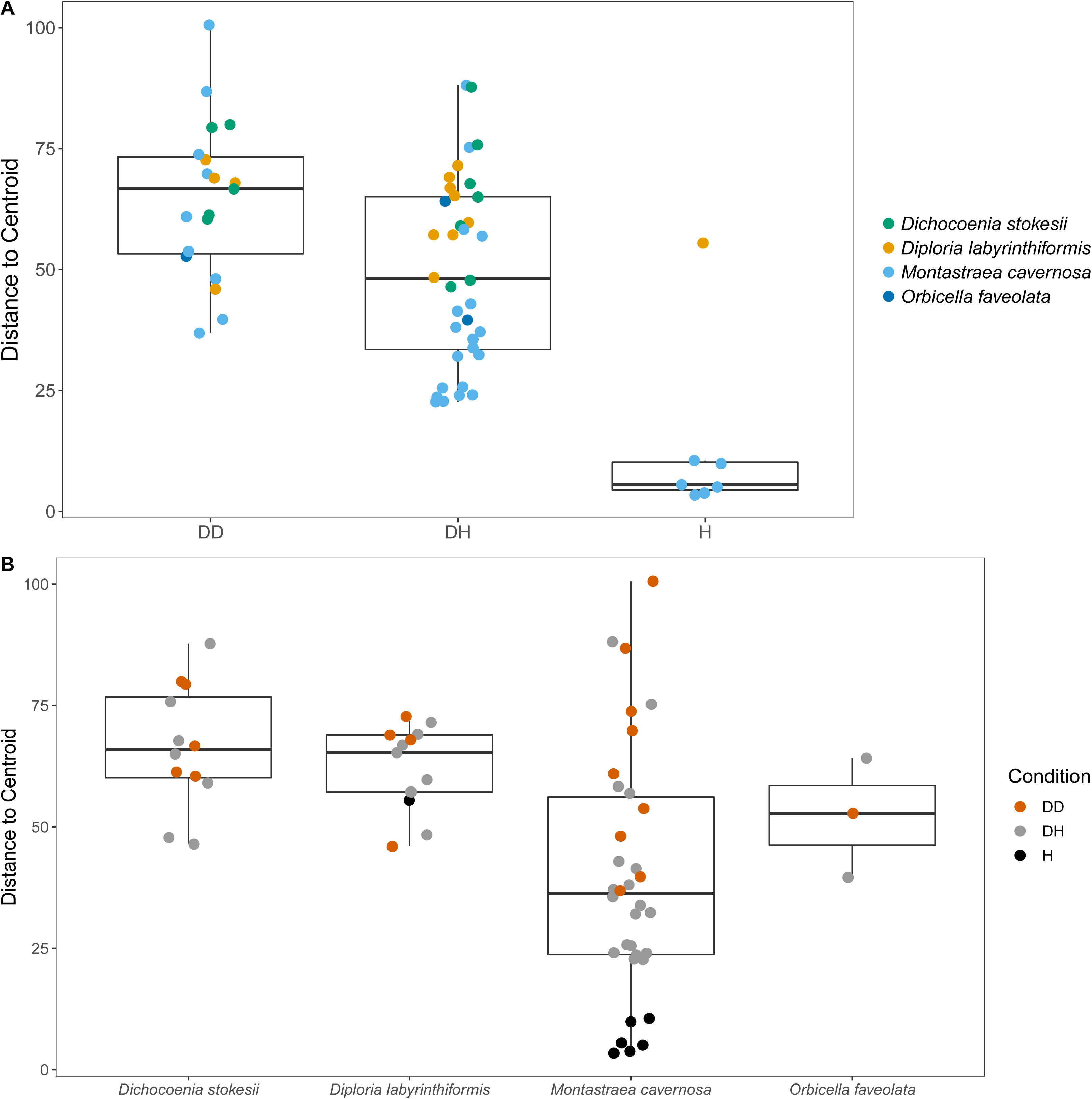
The dispersion of beta diversity shown as the distance to the centroid. Panel (A) compares dispersion of beta diversity in microbial communities from disease lesions (DD), apparently healthy tissue on diseased corals (DH), and apparently healthy neighboring corals with no signs of disease (H), with points colored to indicate coral species. Panel (B) compares dispersion of beta diversity in microbial communities from *Montastraea cavernosa, Orbicella faveolata, Diploria labyrinthiformis*, and *Dichocoenia stokesii*, with points colored to indicate disease condition.

Common taxa detected in all samples of *D. labyrinthiformis* from the middle Keys included Rhodobacteraceae of the genus *Shimia* and Cyanobiaceae, namely *Prochlorococcus* strain MIT9313 and *Synechococcus* strain CC9902 (Figure S2). The healthy neighboring *D. labyrinthiformis* colony and the healthy tissue farthest from the lesion on one of the diseased *D. labyrinthiformis* colonies had high relative abundances of sequences classified as family Simkaniaceae, genus *Candidatus* Fritschea in the Phylum Chlamydiae; members of this group are known as insect endosymbionts (Everett et al., 2005). The healthy colony and healthy tissues also tended to have higher relative abundances of an ASV classified only as Bacteria, that was 99% identical to both a clone library sequence (GenBank Accession KC668983) detected in *Stylophora pistillata* coral from the Red Sea (Bayer et al., 2013) and to a clone (GenBank Accession GQ413901) detected in *Porites cylindrica* in the Philippines (Garren et al., 2009).

Common taxa detected in apparently healthy samples of *D. stokesii* from the middle Keys included an ASV classified only as Bacteria, that was 97% similar to a clone library sequence (GenBank Accession HQ189553) from an estuarine anemone, *Nematostella vectensis*, sampled from Cape Cod (Har et al., 2015). Other common taxa in *D. stokesii* included the Cyanobiaceae strains *Prochlorococcus* MIT9313 and *Synechococcus* CC9902 and an unclassified betaproteobacterial genus of Rhodocyclaceae that was detected in every *D. stokesii* sample (Figure S3). The closest BLASTn matches to this Rhodocyclaceae ASV were 98% similar to sequences recovered from soils. High relative abundances of an ASV that was an exact match to the type species of *Vibrio ishigakensis* from Japanese coral reef seawater (Gao et al., 2016) were detected in the disease lesions of *D. stokesii* colonies as well as in apparently healthy tissues at lower relative abundances.

While microbial community structure changed with sample condition (diseased, apparently healthy tissue on diseased coral, or healthy neighboring coral), the effect size was small (PERMANOVA R^2^ = 0.074, p = 0.001) when compared to the effect of site (PERMANOVA R^2^= 0.154, p = 0.001) or coral species (PERMANOVA R^2^ = 0.223, p = 0.001). Therefore, we looked for taxa that were statistically more abundant (enriched) as determined by DESeq2 analysis in disease lesions within each coral species (Figure S4). In *M. cavernosa*, 119 ASVs were enriched in disease lesions versus healthy tissue on diseased colonies out of 121 ASVs that were detected as differentially abundant. In *D. labyrinthiformis*, 124 of 132 differentially abundant ASVs were enriched in disease lesions. In *D. stokesii*, 69 of 161 differentially abundant ASVs were enriched in disease lesions. Of these, 30 ASVs were enriched in disease lesions of at least two coral species. One of the 30 ASVs, classified as Rhodobacteraceae, was also detected in the PCR blank sample (Table S2). Only five ASVs were enriched in the disease lesions of all three coral species (Table 1, Figure 5). These five ASVs were not detected in any of the control samples (Table S2). The disease-enriched ASVs from all three coral species included an unclassified genus of Flavobacteriales and sequences identified as *Fusibacter* (Clostridiales), *Planktotalea* (Rhodobacterales), *Algicola* (Alteromonadales), and *Vibrio* (Vibrionales).

**Table 1.**
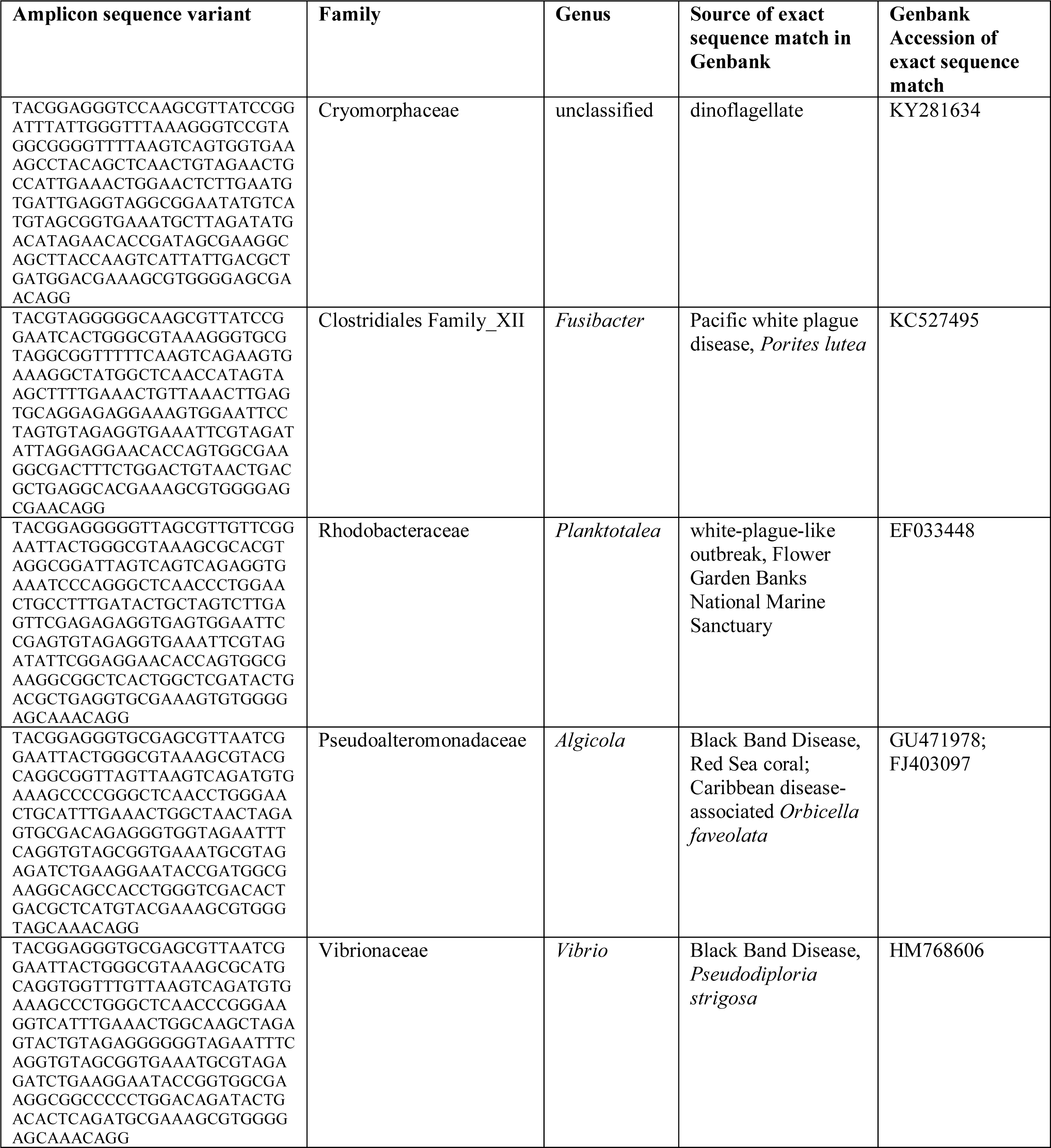
Amplicon sequence variants detected as differentially abundant in stony coral tissue loss disease lesions compared to apparently healthy tissue on diseased corals in *Montastraea cavernosa, Diploria labyrinthiformis*, and *Dichocoenia stokesii*.

**Figure 5.**
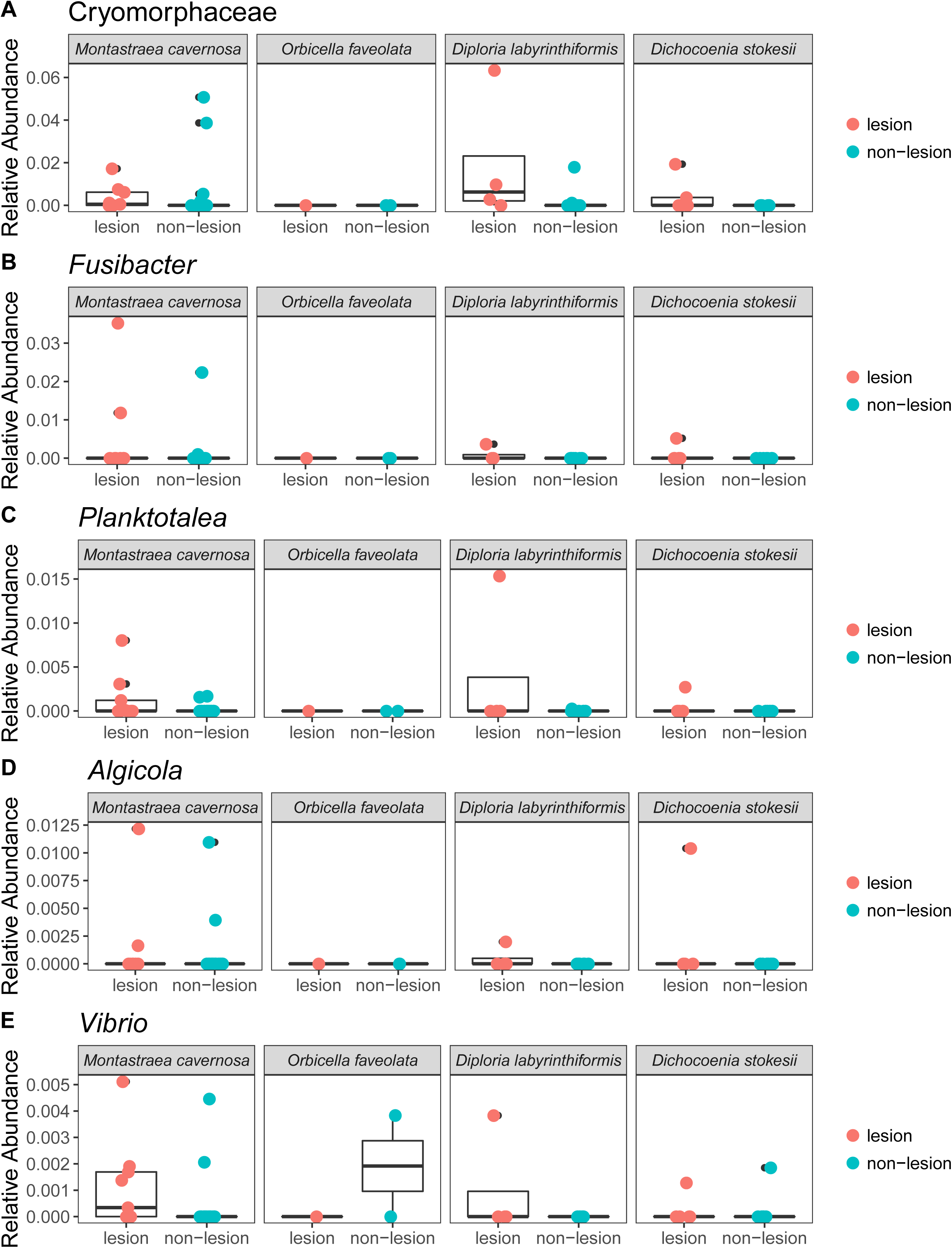
Relative abundance of the three amplicon sequence variants that were differentially abundant in disease lesions versus apparently healthy (non-lesion) tissues of *Montastraea cavernosa* (Mcav), *Orbicella faveolata* (Ofav), *Diploria labyrinthiformis* (Dlab), and *Dichocoenia stokesii* (Dsto).

The Flavobacteriales ASV was identified to family Cryomorphaceae, but not assigned to a genus. This ASV had only one exact BLASTn match to a clone sequence associated with marine dinoflagellates (GenBank Accession KY281634). This Cryomorphaceae ASV was up 6.3% relative abundance in disease lesions of *D. labyrinthiformis* and *D. stokesii* (Figure 5, panel A). The Cryomorphaceae ASV was detected in 10 out of 19 lesion samples, six out of 18 samples of apparently healthy tissue near the disease lesion, and in one out of seven undiseased coral samples. It was not detected in the 18 samples of apparently healthy tissue far from the disease lesion.

The *Fusibacter* ASV was an exact match to a clone library sequence (GenBank Accession KC527495) from a white plague disease infected *Porites lutea* coral from Thailand (Roder et al., 2014) in addition to closely matching (99.6% identical) sequences from Black Band Disease infected corals in the Red Sea (GenBank Accession GU472060, EF089469) (Arotsker et al., 2015). The *Fusibacter* ASV was up to 3.5% relative abundance in the disease lesions of *M. cavernosa* and was detectable at lower levels in disease lesions of *D. labyrinthiformis* and *D. stokesii* (Figure 5, panel B). It was detected in four of 19 lesion samples and two of 18 near lesion samples, but not detected in samples far from the lesion or undiseased corals.

BLASTn searches of the sequences revealed that the *Planktotalea* ASV is a commonly detected strain in marine environments, including in sediments and associated with invertebrates and is also an exact match to an unpublished sequence (GenBank Accession EF033448) associated with an outbreak of white-plague-like disease in the Flower Garden Banks National Marine Sanctuary. The *Planktotalea* ASV was up to 1.5% relative abundance in the disease lesions of *D. labyrinthiformis* and was more abundant in lesions versus non-lesions in *M. cavernosa* and *D. stokesii* (Figure 5, panel C). It was detected in six of 19 lesions and three of 18 near lesion samples, but not detected in samples far from the lesion or undiseased corals.

The *Algicola* ASV was an exact match to a clone library sequence (GenBank Accession GU471978) from a Black Band Disease mat on Red Sea *Favia* sp. (Arotsker et al., 2015) as well as to a clone library sequence (GenBank Accession FJ403097) from an unpublished study of diseased and healthy Caribbean *Orbicella faveolata*. The *Algicola* ASV was up to 1.2% relative abundance in both apparently healthy and diseased tissues of *M. cavernosa* and was more abundant in lesions versus non-lesions in *D. labyrinthiformis* and *D. stokesii* (Figure 5, panel D). It was detected in four of 19 lesions and one sample near the lesion and one sample far from the lesion in *M. cavernosa*.

The *Vibrio* ASV was an exact match to clone library sequences from a Black Band Disease mat on Caribbean *Pseudodiploria strigosa* (GenBank Accession HM768606) (Klaus et al., 2011), from a Black Band Disease mat on Red Sea *Favia* sp. (GenBank Accession GU472063) *(Arotsker et al*., *2015)*, and from white plague disease type II in Caribbean *Orbicella faveolata* (GenBank Accession FJ202106) (Sunagawa et al., 2009). The highest relative abundances of the *Vibrio* ASV were 0.5% in the disease lesion of one *M. cavernosa* colony and was more abundant in lesions versus non-lesions in *D. labyrinthiformis* and *D. stokesii* (Figure 5, panel E). It was detected in seven of 19 lesions and four of 18 near lesion samples, but not detected in samples far from lesions or undiseased corals.

While only five unique ASV sequences were detected at statistically higher abundance in the disease lesions of all three coral species, some groups were consistently detected in higher abundances in disease lesions, albeit with varying ASV sequences. Epsilonbacteraeota, especially Campylobacterales of the genus *Arcobacter*, were more abundant in lesions from all three coral species (Figure S4), including seven ASVs in *M. cavernosa*, six ASVs in *D. labyrinthiformis*, and two ASVs in *D. stokesii*. Only one of these *Arcobacter* ASVs was enriched in more than one coral species (*M. cavernosa* and *D. labyrinthiformis*). In addition, other Campylobacterales were enriched in disease lesions, including *Sulfurospirillum* and *Sulfurovum* in *M. cavernosa, Thiovulum* in *D. labyrinthiformis*, and an Rs-M59_termite_group ASV in *D. stokesii*. Likewise, Patescibacteria were also enriched in disease lesions of all three coral species. Of the 39 ASVs classified as JGI_0000069-P22 in the phylum Patescibacteria, seven were enriched in lesions. Only one ASV classified as JGI_0000069-P22 was enriched in more than one coral species (*D. labyrinthiformis* and *D. stokesii*), which was 96% similar to a sequence (GenBank Accession EU183997) recovered from heat-stressed *Rhopaloeides odorabile* sponge (Webster et al., 2008). In addition, one ASV classified as Gracilibacteria in the phylum Patescibacteria was enriched in lesions on *M. cavernosa* that is an exact match to a clone library sequence (GenBank Accession KC527297) from a white plague disease infected *Pavona duerdeni* coral from Thailand (Roder et al., 2014). One ASV classified as *Fusibacter* (Clostridiales) enriched in lesions from both *M. cavernosa* and *D. labyrinthiformis* was 99.6% identical to the ASV sequence that was differentially abundant in all three coral species. In addition, the *Fusibacter* ASV enriched in lesions from both *M. cavernosa* and *D. labyrinthiformis* was an exact match to three different clone library sequences (GenBank Accessions MH341654, GU472060, EF089469) from Black Band Disease infected corals in the Red Sea (Arotsker et al., 2015; Hadaidi et al., 2018).

In addition to the Flavobacteriales (Cryomorphaceae) ASV enriched in the disease lesions of all three coral species, other Bacteroidetes ASVs were also consistently enriched in disease lesions across all three coral species, including 16 ASVs classified as Bacteroidales, 12 from Chitinophagales, 25 from Cytophagales, and 22 from Flavobacteriales. Of the 16 Bacteroidales ASVs enriched in disease samples, six were exact matches to sequences previously detected in coral diseases, including Black Band Disease in Caribbean *Pseudodiploria strigosa* (GenBank Accession AY497296) (Frias-Lopez et al., 2004), white plague disease in *Porites lutea* from Thailand (GenBank Accession KC527457, KC527469) (Roder et al., 2014), white plague disease type II in Caribbean *Orbicella faveolata* (GenBank Accession FJ202346, FJ202437, FJ202774) (Sunagawa et al., 2009), and Black Band Disease infected corals in the Red Sea (GenBank Accession MH341668, MH341680) (Hadaidi et al., 2018). This included two ASVs with exact matches to sequences from two different disease studies. In addition, one Bacteroidales ASV was a close match (99.6% identical) to sequences (GenBank Accession MH341668, KC527457) from these studies. Three Flavobacteriales ASVs were also matches to sequences previously detected in coral diseases, including an exact match to an unpublished sequence (GenBank Accession EU780277) associated with the disease margin of *Tubinaria mesenterina* and close matches (99.6%) to sequences from Black Band Disease infected *Favia* sp. in the Red Sea (GenBank Accession GU472416, GU471979) (Arotsker et al., 2015).

In addition to the *Planktotalea* (Rhodobacteraceae) enriched in the disease lesions of all three coral species, five additional ASVs classified as Rhodobacteraceae were enriched in the disease lesions of both *M. cavernosa* and *D. labyrinthiformis*, three of which match sequences previously associated with coral disease. Two Rhodobacteraceae ASVs were exact matches to clone library sequences from Black Band Disease (GenBank Accession DQ446109, MH341655) (Hadaidi et al., 2018; Sekar et al., 2008) and one Rhodobacteraceae ASV was an exact match to a clone library sequence from white plague disease type II (GenBank Accession FJ203176) (Sunagawa et al., 2009).

In addition to the *Vibrio* ASV enriched in the disease lesions of all three coral species, one additional *Vibrio* ASV was enriched in the disease lesions of *M. cavernosa* and *D. labyrinthiformis* that was an exact match to a sequence (GenBank Accession MK168657) identified as *V. coralliilyticus* strain 2214. While this particular strain is not known to be pathogenic, other strains within this species are known pathogens of coral and shellfish (Ushijima et al., 2014, 2018). It should be noted that this *Vibrio* ASV is 253 bp long and is an equally good match to dozens of other *Vibrio* species and strains, making an exact identification of *Vibrio* strain impossible with the V4 region of the 16S small subunit ribosomal gene.

## DISCUSSION

Consistent with other microbial investigations of coral diseases, we found a shift in the microorganisms present in stony coral tissue loss disease (SCTLD) disease lesions versus apparently healthy coral tissue. The disruption, or dysbiosis, of the normal, healthy microbiota, which in corals is often dominated by a small number of bacterial groups, is often paralleled by an increase in microbial diversity. This increased diversity is often stochastic, so that each altered microbiome is unique (Zaneveld et al., 2017), while the original healthy microbial community has a stable composition of lower diversity. In the current study, this pattern held true for *M. cavernosa*, from which we sampled several undiseased colonies. The undiseased colonies as well as apparently healthy tissue on diseased colonies sampled in summer from *M. cavernosa* were dominated by the gammaproteobacterial genera *Halomonas* and *Shewanella*. In contrast, the apparently healthy tissue on diseased colonies sampled in winter were much more variable, like the microbiota associated with lesions. In *D. labyrinthiformis* and *D. stokesii*, both of which were sampled in winter, apparently healthy tissue on diseased colonies as well as disease lesions hosted diverse microbial communities, and no difference was seen in the dispersion of beta diversity between healthy and diseased tissue on diseased colonies. This suggests that when sampling when the disease appears to be more active and when sampling from more susceptible coral species (Florida Keys National Marine Sanctuary, 2018), the microbiome of the entire colony is already disrupted by SCTLD. The December sampling also followed extreme environmental conditions and changes in salinity, turbidity, and sedimentation caused by Hurricane Irma in September 2017, which could have also affected microbiome composition of all the corals sampled in winter.

A single potential pathogen was not identified; however, five unique ASV sequences were enriched in the SCTLD lesions of three different coral species, and all but one of these sequences were exact matches to sequences previously associated with coral disease. These sequences were classified as Flavobacteriales, Clostridiales, Rhodobacterales, Alteromonadales, and Vibrionales, groups which have all been consistently detected in coral diseases, as well as in apparently healthy coral tissues. Previous investigations of white plague disease type II in Caribbean *O. faveolata* also found an enrichment of the same five bacterial orders (Sunagawa et al., 2009). The *Fusibacter* ASV sequence enriched in the lesions of all three coral species is an exact match to a sequence from coral infected with white plague disease in the Pacific (Roder et al., 2014). The unique *Planktotalea* sequence (Rhodobacterales) was an exact match to an unpublished sequence previously detected in a white-plague-type outbreak in the Gulf of Mexico, and the *Vibrio* ASV sequence enriched in the lesions of all three coral species is an exact match to a sequence enriched in the white plague disease type II lesions from the Sunagawa *et al*. study. However, the enrichment of these groups is not confined to “white syndrome” coral diseases. Previous work has also identified the enrichment of Flavobacteriales, Clostridiales, Rhodobacterales, Alteromonadales, and Vibrionales associated with Black Band Disease mats (Miller and Richardson, 2011). Indeed, ASV sequences enriched in disease lesions of SCTLD were also exact matches to sequences enriched in Black Band Disease from geographically disparate tropical regions.

Collectively, the presence of similar microbes in several coral diseases from around the world can be interpreted in multiple ways. These microbes may be the pathogens that initiate coral disease, however, their presence in multiple, visually distinct coral diseases suggests that this may not be the case. In addition to primary pathogens, these microbes may be opportunistic or secondary pathogens that infect immunocompromised or otherwise vulnerable corals (Lesser et al., 2007), such as those stressed by elevated water temperatures or sedimentation. The current outbreak was preceded by both higher sea surface temperatures in the summer and winter of 2014 (Manzello, 2015) and higher sedimentation on corals due to dredging operations between 2013 and 2015 in the channel at the Port of Miami (Miller et al., 2016). Coral microbiome diversity increases with environmental stresses such as climate change, water pollution, and overfishing and is accompanied by an increase of bacteria classified as *Vibrionales, Flavobacteriales, Rhodobacterales, Alteromonadales, Rhizobiales, Rhodospirillales*, and *Desulfovibrionales* (McDevitt-Irwin et al., 2017). Here, we detected an enrichment of the first four of these bacterial orders in the disease lesions of SCTLD. Organisms that are enriched in the disease state may be resident bacteria that are already present in healthy corals and grow in response to changes in the host during disease progression. Recent experimental infections of *Acropora cervicornis* with white plague disease revealed both colonizers present in the diseased coral tissue that transferred to and increased in newly infected corals as well as responders that were present in healthy tissue and increased during disease progression in newly infected corals (Gignoux-Wolfsohn et al., 2017).

Alternatively, the disease-enriched microbes may be saprophytic colonizers that increase in numbers as decaying coral tissue fuels their growth (Egan and Gardiner, 2016). In particular, many of the enriched groups detected here are associated with the anoxic conditions that accompany tissue decay, such as the Epsilonbacteraeota (Campbell et al., 2006), Clostridiales such as *Fusibacter* (Ravot et al., 1999), *and* Patescibacteria. The superphylum Patescibacteria is comprised of three environmentally widespread phyla that to-date have no cultured representatives, but metagenomic analyses have revealed that Patescibacteria lack TCA cycle genes and therefore must be strict anaerobic fermenters (Wrighton et al., 2012) and that they tend to have reduced genomes, characteristic of parasitic or symbiotic microbes (Sánchez-Osuna et al., 2017). Our study may be the first to specifically detect this relatively newly named superphylum (Rinke et al., 2013) as present in coral disease lesions; however, one Patescibacteria ASV sequence detected here matched a clone library sequence from white plague disease infected *Pavona duerdeni* coral from Thailand (Roder et al., 2014), suggesting that these cryptic yet environmentally widespread taxa may also be common in coral disease.

The enrichment of disease-associated bacteria in the lesions of corals with SCTLD is not definitive proof that the pathogen is bacterial. However, disease progression in laboratory and field trials appears to slow or stop with the application of antibiotics (Florida Keys National Marine Sanctuary, 2018), strongly suggesting that bacteria are involved with disease progression. The isolation and culturing of a bacterial pathogen(s) would be invaluable to the development of diagnostic methods and targeted mitigation efforts. Therefore, attempts at isolation of the potential pathogen and establishing new infections with isolates is ongoing by our group. In addition, the pathogen or pathogens responsible for SCTLD may be viral or eukaryotic, which may only be uncovered through techniques such as shotgun metagenomics. Metagenomic analysis of samples from corals impacted by SCTLD may provide additional information regarding the nature of the pathogen, whether bacterial, viral, or eukaryotic, as well as the mechanisms used by the pathogen(s) to infect and kill corals.

## ACKNOWLEDGMENTS

The authors thank Broward County Environmental Protection and Growth Management Department, particularly Angel Rovira and Ken Banks, Keys Marine Laboratory (Cindy Lewis, Bill Ferrell) and Audrey Looby for coordination and assistance in field collections. Field collections were authorized under Florida Fish and Wildlife Conservation Commission Special Activity Licenses SAL-16-1702A-SRP and SAL-17-1702-SRP and Florida Keys National Marine Sanctuary permit # FKNMS-2017-128. This work was supported by an NSF Rapid Grant (IOS-1728002) to VP, CH, GA, and JM and an EPA Wetlands Program Grant (CD00D66917) to JM, CH, and VP. This paper is Smithsonian Marine Station publication number XXXX.

## AUTHOR CONTRIBUTIONS STATEMENT

Experimental design was determined collectively by JM, GA, CH, BU, and VP. Field collections were conducted by JM and VP. Molecular lab work and analysis was performed by JM and JC. JM drafted the manuscript and all authors contributed to interpretations and revisions.

## CONTRIBUTION TO THE FIELD STATEMENT

The Caribbean is known as a hotspot for coral diseases that have dramatically reduced populations of reef-building corals. Currently, half of the coral species in the Florida Reef Tract are impacted by an ongoing outbreak of stony coral tissue loss disease that was first detected in late 2014. This work presents the first characterization of the changes in microbiological communities associated with the disease in four stony coral species. Disease lesions consistently supported microbial communities enriched with bacterial taxa that have previously been detected in coral diseases, including white-plague-type diseases and Black Band Disease. Enriched taxa in disease lesions may be causing the primary infection or may be secondary infections. Additional studies are needed to determine the roles of bacteria enriched in diseased tissue and to evaluate the potential for non-bacterial pathogens such as viruses.

## Supplemental Information

Figure S1. Relative abundance of amplicon sequence variants, colored by Family, in undiseased neighboring corals, apparently healthy tissue far from the disease lesion, apparently healthy tissue near the disease lesion, and disease lesions in *Montastraea cavernosa* with stony coral tissue loss disease. Samples with names beginning with PB were collected in July 2017 and samples with names beginning with a number were collected December 2017.

Figure S2. Relative abundance of amplicon sequence variants, colored by Family, in undiseased neighboring corals, apparently healthy tissue far from the disease lesion, apparently healthy tissue near the disease lesion, and disease lesions in *Diploria labyrinthiformis* with stony coral tissue loss disease.

Figure S3. Relative abundance of amplicon sequence variants, colored by Family, in apparently healthy tissue far from the disease lesion, apparently healthy tissue near the disease lesion, and disease lesions in *Dichocoenia stokesii* with stony coral tissue loss disease.

Figure S4. Differentially abundant amplicon sequence variants (ASVs) in stony coral tissue loss disease. Positive log2FoldChange values correspond to ASVs that are more abundant in disease lesions compared to apparently healthy tissue on diseased corals in *Montastraea cavernosa*.

Figure S5. Differentially abundant amplicon sequence variants (ASVs) in stony coral tissue loss disease. Positive log2FoldChange values correspond to ASVs that are more abundant in disease lesions compared to apparently healthy tissue on diseased corals in *Diploria labyrinthiformis*.

Figure S6. Differentially abundant amplicon sequence variants (ASVs) in stony coral tissue loss disease. Positive log2FoldChange values correspond to ASVs that are more abundant in disease lesions compared to apparently healthy tissue on diseased corals in *Dichocoenia stokesii*.

Table S1. Sample metadata for microbial communities collected from diseased and apparently healthy tissue associated with stony coral tissue loss disease. Condition DD indicates disease lesion, DH indicates apparently healthy tissue on diseased colonies (and is further parsed as near or far from the lesion), H indicates apparently healthy neighboring corals. Up to 3 samples were collected per coral colony, as indicated by the colony name. Raw reads (with adapters removed) are deposited in NCBI under Bioproject Accession # PRJNA521988.

Table S2. Amplicon sequence variants detected in three control samples. Blank 1 (B1) and Blank 2 (B2) were no template controls from DNA extraction kit through sequencing. Blank 3 (B3) was a no template PCR control that was cleaned and sequenced.

