## Supplemental tables and figures for "Microbial community shifts associated with the ongoing stony coral tissue loss disease outbreak on the Florida Reef Tract"

Table S1. Sample metadata for microbial communities collected from diseased and apparently healthy tissue associated with stony coral tissue loss disease. Condition DD indicates disease lesion, DH indicates apparently healthy tissue on diseased colonies (and is further parsed as near or far from the lesion), H indicates apparently healthy neighboring corals. Up to 3 samples were collected per coral colony, as indicated by the colony name. Raw reads (with adapters removed) are deposited in NCBI under Bioproject Accession # PRJNA521988.

| Sample name | Site | Sample date | Coral | Colony | Condition | Near_Far | raw reads | reads used | NCBI Biosample Accession # |
|---|---|---|---|---|---|---|---|---|---|
| 10FTL | Ft. Lauderdale | Dec-17 | Montastraea cavernosa | mcav1 | DD | lesion | 163206 | 97735 | SAMN10964620 |
| 11FTL | Ft. Lauderdale | Dec-17 | Montastraea cavernosa | mcav1 | DH | near | 16564 | 11589 | SAMN10964621 |
| 12FTL | Ft. Lauderdale | Dec-17 | Montastraea cavernosa | mcav1 | DH | far | 32531 | 7695 | SAMN10964622 |
| 13FTL | Ft. Lauderdale | Dec-17 | Orbicella faveolata | ofav1 | DH | far | 77087 | 31860 | SAMN10964623 |
| 14FTL | Ft. Lauderdale | Dec-17 | Orbicella faveolata | ofav1 | DH | near | 52715 | 8627 | SAMN10964624 |
| 15FTL | Ft. Lauderdale | Dec-17 | Orbicella faveolata | ofav1 | DD | lesion | 59267 | 3288 | SAMN10964625 |
| 1FTL | Ft. Lauderdale | Dec-17 | Montastraea cavernosa | mcav2 | DD | lesion | 138377 | 26167 | SAMN10964626 |
| 2FTL | Ft. Lauderdale | Dec-17 | Montastraea cavernosa | mcav2 | DH | near | 126210 | 43396 | SAMN10964627 |
| 3FTL | Ft. Lauderdale | Dec-17 | Montastraea cavernosa | mcav2 | DH | far | 100050 | 19103 | SAMN10964628 |
| 4FTL | Ft. Lauderdale | Dec-17 | Montastraea cavernosa | mcav3 | DD | lesion | 9678 | 3513 | SAMN10964629 |
| 5FTL | Ft. Lauderdale | Dec-17 | Montastraea cavernosa | mcav3 | DH | near | 94544 | 15900 | SAMN10964630 |
| 6FTL | Ft. Lauderdale | Dec-17 | Montastraea cavernosa | mcav3 | DH | far | 37793 | 13040 | SAMN10964631 |
| 7FTL | Ft. Lauderdale | Dec-17 | Montastraea cavernosa | mcav4 | DD | lesion | 122318 | 77995 | SAMN10964632 |
| 8FTL | Ft. Lauderdale | Dec-17 | Montastraea cavernosa | mcav4 | DH | near | 151987 | 72875 | SAMN10964633 |
| 9FTL | Ft. Lauderdale | Dec-17 | Montastraea cavernosa | mcav4 | DH | far | 72044 | 7286 | SAMN10964634 |
| Dl10A | Long Key Bridge | Dec-17 | Diploria labyrinthiformis | dl10 | DD | lesion | 175233 | 59052 | SAMN10964635 |
| Dl10B | Long Key Bridge | Dec-17 | Diploria labyrinthiformis | dl10 | DH | near | 145483 | 49168 | SAMN10964636 |
| Dl10C | Long Key Bridge | Dec-17 | Diploria labyrinthiformis | dl10 | H | undiseased | 162793 | 95389 | SAMN10964637 |
| Dl6A | Long Key Bridge | Dec-17 | Diploria labyrinthiformis | dl6 | DD | lesion | 144530 | 119587 | SAMN10964638 |
| Dl6B | Long Key Bridge | Dec-17 | Diploria labyrinthiformis | dl6 | DH | near | 107287 | 68671 | SAMN10964639 |
| Dl6C | Long Key Bridge | Dec-17 | Diploria labyrinthiformis | dl6 | DH | far | 91447 | 44119 | SAMN10964640 |
| Dl7C | Long Key Bridge | Dec-17 | Diploria labyrinthiformis | dl7 | DH | far | 111669 | 24228 | SAMN10964641 |
| Dl8A | Long Key Bridge | Dec-17 | Diploria labyrinthiformis | dl8 | DD | lesion | 56113 | 20045 | SAMN10964642 |
| Dl8B | Long Key Bridge | Dec-17 | Diploria labyrinthiformis | dl8 | DH | near | 85465 | 21961 | SAMN10964643 |
| Dl8C | Long Key Bridge | Dec-17 | Diploria labyrinthiformis | dl8 | DH | far | 56643 | 2358 | SAMN10964644 |
| Dl9A | Long Key Bridge | Dec-17 | Diploria labyrinthiformis | dl9 | DD | lesion | 159012 | 85597 | SAMN10964645 |
| Dl9B | Long Key Bridge | Dec-17 | Diploria labyrinthiformis | dl9 | DH | near | 189498 | 40454 | SAMN10964646 |
| Dl9C | Long Key Bridge | Dec-17 | Diploria labyrinthiformis | dl9 | DH | far | 244958 | 85801 | SAMN10964647 |
| Ds1A | Long Key Bridge | Dec-17 | Dichocoenia stokesii | ds1 | DD | lesion | 159132 | 107824 | SAMN10964648 |
| Ds1B | Long Key Bridge | Dec-17 | Dichocoenia stokesii | ds1 | DH | near | 72383 | 35677 | SAMN10964649 |
| Ds1C | Long Key Bridge | Dec-17 | Dichocoenia stokesii | ds1 | DH | far | 100124 | 57394 | SAMN10964650 |

| | | | | | | | | | |
|---|---|---|---|---|---|---|---|---|---|
| Ds2A | Long Key Bridge | Dec-17 | Dichocoenia stokesii | ds2 | DD | lesion | 71721 | 36451 | SAMN10964651 |
| Ds2B | Long Key Bridge | Dec-17 | Dichocoenia stokesii | ds2 | DH | near | 64309 | 15308 | SAMN10964652 |
| Ds2C | Long Key Bridge | Dec-17 | Dichocoenia stokesii | ds2 | DH | far | 51361 | 17476 | SAMN10964653 |
| Ds3A | Long Key Bridge | Dec-17 | Dichocoenia stokesii | ds0 | DD | lesion | 164114 | 93611 | SAMN10964654 |
| Ds3B | Long Key Bridge | Dec-17 | Dichocoenia stokesii | ds3 | DD | lesion | 137638 | 46421 | SAMN10964655 |
| Ds3C | Long Key Bridge | Dec-17 | Dichocoenia stokesii | ds3 | DH | far | 156119 | 85671 | SAMN10964656 |
| Ds4A | Long Key Bridge | Dec-17 | Dichocoenia stokesii | ds4 | DD | lesion | 111770 | 38470 | SAMN10964657 |
| Ds4B | Long Key Bridge | Dec-17 | Dichocoenia stokesii | ds4 | DH | near | 102816 | 26344 | SAMN10964658 |
| Ds4C | Long Key Bridge | Dec-17 | Dichocoenia stokesii | ds4 | DH | far | 122982 | 40126 | SAMN10964659 |
| PBa | Ft. Lauderdale | Jul-17 | Montastraea cavernosa | a | H | undiseased | 8605 | 3741 | SAMN10964660 |
| PBA1 | Ft. Lauderdale | Jul-17 | Montastraea cavernosa | A | DH | far | 28274 | 14874 | SAMN10964661 |
| PBA2 | Ft. Lauderdale | Jul-17 | Montastraea cavernosa | A | DH | near | 35187 | 9102 | SAMN10964662 |
| PBA3 | Ft. Lauderdale | Jul-17 | Montastraea cavernosa | A | DD | lesion | 185193 | 109492 | SAMN10964663 |
| PBA4 | Ft. Lauderdale | Jul-17 | Montastraea cavernosa | A | DH | near | 128398 | 44379 | SAMN10964664 |
| PBb | Ft. Lauderdale | Jul-17 | Montastraea cavernosa | b | H | undiseased | 6863 | 5478 | SAMN10964665 |
| PBB1 | Ft. Lauderdale | Jul-17 | Montastraea cavernosa | B | DH | far | 68874 | 44928 | SAMN10964666 |
| PBB2 | Ft. Lauderdale | Jul-17 | Montastraea cavernosa | B | DH | near | 79613 | 44691 | SAMN10964667 |
| PBB3 | Ft. Lauderdale | Jul-17 | Montastraea cavernosa | B | DD | lesion | 338067 | 233271 | SAMN10964668 |
| PBc | Ft. Lauderdale | Jul-17 | Montastraea cavernosa | c | H | undiseased | 17609 | 10740 | SAMN10964669 |
| PBC1 | Ft. Lauderdale | Jul-17 | Montastraea cavernosa | C | DH | far | 41706 | 27151 | SAMN10964670 |
| PBC2 | Ft. Lauderdale | Jul-17 | Montastraea cavernosa | C | DH | near | 90670 | 37158 | SAMN10964671 |
| PBC3 | Ft. Lauderdale | Jul-17 | Montastraea cavernosa | C | DD | lesion | 79129 | 67744 | SAMN10964672 |
| PBC4 | Ft. Lauderdale | Jul-17 | Montastraea cavernosa | c2 | H | undiseased | 3860 | 3116 | SAMN10964673 |
| PBd | Ft. Lauderdale | Jul-17 | Montastraea cavernosa | d | H | undiseased | 11947 | 9613 | SAMN10964674 |
| PBD1 | Ft. Lauderdale | Jul-17 | Montastraea cavernosa | D | DH | far | 60338 | 37327 | SAMN10964675 |
| PBD2 | Ft. Lauderdale | Jul-17 | Montastraea cavernosa | D | DH | near | 30105 | 9809 | SAMN10964676 |
| PBD3 | Ft. Lauderdale | Jul-17 | Montastraea cavernosa | D | DD | lesion | 12906 | 10555 | SAMN10964677 |
| PBF1 | Ft. Lauderdale | Jul-17 | Montastraea cavernosa | f | H | undiseased | 9843 | 5568 | SAMN10964678 |
| PBF2 | Ft. Lauderdale | Jul-17 | Montastraea cavernosa | F | DH | far | 10623 | 8384 | SAMN10964679 |
| PBF3 | Ft. Lauderdale | Jul-17 | Montastraea cavernosa | F | DH | near | 59953 | 26104 | SAMN10964680 |
| PBF4 | Ft. Lauderdale | Jul-17 | Montastraea cavernosa | F | DD | lesion | 175522 | 118210 | SAMN10964681 |
| B1 | extraction blank | n/a | n/a | n/a | n/a | n/a | 1114 | 258 | SAMN10964682 |
| B2 | extraction blank | n/a | n/a | n/a | n/a | n/a | 97960 | 18348 | SAMN10964683 |
| B3 | PCR blank | n/a | n/a | n/a | n/a | n/a | 54582 | 11827 | SAMN10964684 |

Table S2. Amplicon sequence variants detected in 3 control samples. Blank 1 (B1) and Blank 2 (B2) were no template controls from DNA extraction kit through sequencing. Blank 3 (B3) was a no template PCR control that was cleaned and sequenced.

| Kingdom | Phylum | Class | Order | Family | Genus | blank total | B1 | B2 | B3 | Amplicon Sequence Variant |
|---|---|---|---|---|---|---|---|---|---|---|
| Bacteria | Acidobacteria | Acidobacteriia | Solibacterales | Solibacteraceae_(Subgroup_3) | Bryobacter | 157 | 0 | 0 | 157 | TACGTAGGCAGCGAGCGTTGTTCGGAATTACTGGGCGTAAAGAGTGTGTAGGCGGTGCTCTAAGTTCGGTGTGAAATCTCCCGGCTTAACTGGGAGGGTGCGCCGGAAACTGGAGTGCTCGAGCGTGGGAGAGGAAAGCGGAATTCCTGGTGTAGCGGTGAAATGCGTAGATATCAGGAGGAACACCTGCGGTGTAGACGGCTTTCTGGACCATTGCTGACGCTGAGACACGAAAGCGTGGGTAGCAAACAGG |
| Bacteria | Acidobacteria | Acidobacteriia | Solibacterales | Solibacteraceae_(Subgroup_3) | Solibacteraceae_(Subgroup_3) | 137 | 0 | 0 | 137 | TACGTAGGCAGCGAGCGTTGTTCGGAATTACTGGGCGTAAAGAGTGTGTAGGCGGTGCTCTAAGTTCGGTGTGAAATCTCCCGGCTTAACTGGGAGGGTGCGCCGGAAACTGGAGTGCTCGAGCGTGGGAGAGGAAAGCGGAATTCCTGGTGTAGCGGTGAAATGCGTAGATATGTGGAAGAACACCCGTGGCGAAGGCGGCATCCTGGACCGGTATTGACGCTGATGCGCGAAAGCCAGGGGAGCAAACGGG |
| Bacteria | Actinobacteria | Actinobacteria | Corynebacteriales | Nocardiaceae | Gordonia | 114 | 0 | 0 | 114 | TACGTAGGGTGCGAGCGTTGTCCGGAATTACTGGGCGTAAAGAGCTCGTAGGCGGTTTGTCGCGTCGTCTGTGAAATTCTGCAGCTTAACTGCAGGCGTGCAGGCGATACGGGCAGACTTGAGTACTACAGGGGAGACTGGAATTCCTGGTGTAGCGGTGAAATGCGCAGATATCAGGAGGAACACCGGTGGCGAAGGCGGGTCTCTGGGTAGTAACTGACGCTGAGGAGCGAAAGCGTGGGTAGCGAACAGG |
| Bacteria | Actinobacteria | Actinobacteria | Corynebacteriales | Tsukamurellaceae | Tsukamurella | 32 | 0 | 0 | 32 | TACGTAGGGTGCGAGCGTTGTCCGGATTCACTGGGCGTAAAGAGCTCGTAGGCGGTTTGTCACGTCGTCTGTGAAAACCCGAGGCTTAACCTCGGGCCTGCAGGCGATACGGGCAGACTTGAGTACTGTAGGGGAGACTGGAATTCCTGGTGTAGCGGTGGAATGCGCAGATATCAGGAGGAACACCGGTGGCGAAGGCGGGTCTCTGGGCAGTAACTGACGCTGAGGAGCGAAAGCGTGGGTAGCGAACAGG |
| Bacteria | Actinobacteria | Actinobacteria | Micrococcales | Microbacteriaceae | Microbacteriaceae | 261 | 0 | 261 | 0 | TACGTAGGGTGCAAGCGTTGTCCGGAATTATTGGGCGTAAAGAGCTCGTAGGCGGTTTGTCACGTCTGCTGTGAAATTCCGAGGCTCAACCTCGGACTTGCAGTGGGTACGGGCAGGCTAGAGTGCGGTAGGGGAGATGGGAATTCCTGGTGTAGCGGTGGAATGCGTAGATATGCGGAGGAACACCGATGGCGAAGGCAATCCCCTGGGCCTGTACTGACGCTCATGCACGAAAGCGTGGGGAGCAAACAGG |
| Bacteria | Actinobacteria | Actinobacteria | Micrococcales | Microbacteriaceae | Parafrigoribacterium | 352 | 0 | 0 | 352 | TACGTAGGGTGCAAGCGTTATCCGGAATTATTGGGCGTAAAGAGCTCGTAGGCGGTTTGTCGCGTCTGCTGTGAAAACTGGAGGCTCAACCTCCAGCCTGCAGTGGGTACGGGCAGACTAGAGTGCGGTAGGGGAGATTGGAATTCCTGGTGTAGCGGTGGAATGCGCAGATATCAGGAGGAACACCAATGGCGAAGGCAGATCTCTGGGCCGTAACTGACGCTGAGGAGCGAAAGCATGGGGAGCGAACAGG |
| Bacteria | Actinobacteria | Actinobacteria | Micrococcales | Micrococcaceae | Renibacterium | 270 | 15 | 0 | 255 | TACGTAGGGTGCAAGCGTTATCCGGAATTATTGGGCGTAAAGAGCTCGTAGGCGGTTTGTCGCGTCTGTCGTGAAAGTCCGGGGCTCAACTCCGGATCTGCGGTGGGTACGGGCAGACTAGAGTGATGTAGGGGAGACTGGAATTCCTGGTGTAGCGGTGGAATGCGCAGATATCAGGAGGAACACCGATGGCGAAGGCAGGTCTCTGGGCATTAACTGACGCTGAGGAGCGAAAGCATGGGGAGCGAACAGG |
| Bacteria | Actinobacteria | Actinobacteria | Micrococcales | Micrococcaceae | Renibacterium | 35 | 0 | 35 | 0 | TACGTAGGGTGCAAGCGTTATCCGGAATTATTGGGCGTAAAGAGCTCGTAGGCGGTTTGTCGCGTCTGTCGTGAAAGTCCGGGGCTCAACTCCGGATCTGCGGTGGGTACGGGCAGACTAGAGTGATGTAGGGGAGACTGGAATTCATGGTGTAGCGGTGGAATGCGCAGATATCAGGAGGAACACCGATGGCGAAGGCAGGTCTCTGGGCATTAACTGACGCTGAGGAGCGAAAGCATGGGGAGCGAACAGG |
| Bacteria | Actinobacteria | Actinobacteria | Pseudonocardiales | Pseudonocardiaceae | Actinomycetospora | 96 | 0 | 0 | 96 | TACGTAGGGTGCGAGCGTTGTCCGGAATTATTGGGCGTAAAGAGCTCGTAGGCGGTGTGTCACGTCGGCCGTGAAAACCTGGGGCTCAACTCTGGGCGTGCGGTCGATACGGGCATCACTTGAGTTCGGCAGGGGAGACTGGAATTCATGGTGTAGCGGTGAAATGCGCAGATATCAGGAGGAACACCGGTGGCGAAGGCGGGTCTCTGGGCCGATACTGACGCTGAGGAGCGAAAGCATGGGTAGCGAACAGG |
| Bacteria | Actinobacteria | Actinobacteria | Pseudonocardiales | Pseudonocardiaceae | Pseudonocardia | 27 | 0 | 0 | 27 | CACGTAGGGCGCGAGCGTTGTCCGGAATTATTGGGCGTAAAGAGCTCGTAGGCGGTGTGTCGCGTCGGCCGTGAAAACCTGCAGCTTAACTGTGGGCTTGCGGTCGATACGGGCATCACTGGAGTTCGGCAGGGGAGACTGGAATTCCTGGTGTAGCGGTGAAATGCGCAGATATCAGGAGGAACACCGGTGGCGAAGGCGGGTCTCTGGGCCGATACTGACGCTGAGGAGCGAAAGCGTGGGGAGCGAACAGG |
| Bacteria | Actinobacteria | Actinobacteria | Streptomycetales | Streptomycetaceae | Allostreptomyces | 377 | 0 | 377 | 0 | TACGTAGGGTGCGAGCGTTGTCCGGAATTATTGGACGTAAAGAGCTCGTAGGCGGCTTGTCGCGTCGGATGTGAAAGCCCGGGGCTTAACTCCGGGTCTGCATTCGATACGGGCAGGCTAGAGTTCGGTAGGGGAGATCGGAATTCCTGGTGTAGCGGTGAAATGCGCAGATATCAGGAGGAACACCGGTGGCGAAGGCAGCACTCTGGGACGTGACTGACGCTGAGACGCGAAAGCGTGGGGAGCAAACAGG |
| Bacteria | Actinobacteria | Actinobacteria | Streptomycetales | Streptomycetaceae | Streptomyces | 150 | 0 | 0 | 150 | TACGTAGGGCGCAAGCGTTGTCCGGAATTATTGGGCGTAAAGAGCTCGTAGGCGGCTTGTCACGTCGGACGTGAAAGCTTGGGGCTTAACTCCAAGTCTGCGTTCGATACGGGCTGGCTAGAGTGTGGTAGGGGAGATCGGAATTCCTGGTGTAGCGATGAAATGCGCAGATATCAGGAGGAACACCGGTGGCGAAGGCGGATCTCTGGGCCATTACTGACGCTGAGGAGCGAAAGCGTGGGGAGCGAACAGG |
| Bacteria | Actinobacteria | Actinobacteria | Streptomycetales | Streptomycetaceae | Streptomyces | 130 | 0 | 0 | 130 | TACGTAGGGCGCAAGCGTTGTCCGGAATTATTGGGCGTAAAGAGCTCGTAGGCGGCTTGTCACGTCGGACGTGAAAGCCCGGGGCTTAACCCCGGGTCTGCAGTCGATACGGGCAGGCTAGAGTTCGGTAGGGGAGATCGGAATTCCTGGTGTAGCGGTGAAATGCGCAGATATCAGGAGGAACACCGGTGGCGAAGGCGGATCTCTGGGCCGATACTGACGCTGAGGAGCGAAAGCGTGGGGAGCGAACAGG |
| Bacteria | Actinobacteria | Actinobacteria | Streptomycetales | Streptomycetaceae | Streptomycetaceae | 1304 | 0 | 445 | 859 | TACGTAGGGCGCAAGCGTTGTCCGGAATTATTGGGCGTAAAGAGCTCGTAGGCGGCTTGTCACGTCGGACGTGAAAGCTTGGGGCTTAACTCCAAGTCTGCGTTCGATACGGGCTGGCTAGAGTGTGGTAGGGGAGATCGGAATTCCTGGTGTAGCGGTGAAATGCGCAGATATCAGGAGGAACACCGGTGGCGAAGGCGGATCTCTGGGCCATTACTGACGCTGAGGAGCGAAAGCGTGGGGAGCGAACAGG |
| Bacteria | Actinobacteria | Actinobacteria | Streptomycetales | Streptomycetaceae | Streptomycetaceae | 179 | 0 | 179 | 0 | TACGTAGGGCGCAAGCGTTGTCCGGAATTATTGGGCGTAAAGAGCTCGTAGGCGGCTTGTCACGTCGGGTGTGAAAGCCCGGGGCTTAACCCCGGGTCTGCATTCGATACGGGCTGGCTAGAGTGTGGTAGGGGAGATCGGAATTCCTGGTGTAGCGGTGAAATGCGCAGATATCGGGAGGAACACCAGTGGCGAAGGCGGCTCACTGGACCACAACTGACGCTGAGGCGCGAAAGCTAGGGGAGCAAACAGG |

| | | | | | | | | | | |
|---|---|---|---|---|---|---|---|---|---|---|
| Bacteria | Actinobacteria | Actinobacteria | Streptosporangiales | Streptosporangiaceae | Acrocarpospora | 46 | 0 | 0 | 46 | TACGTAGGGCGCAAGCGTTGTCCGGAATTATTGGGCGTAAAGAGCTCGTAGGCGGCCTGTTGCGTCTG<br>CCGTGAAAGCCCGTGGCTTAACTGCGGGTCTGCGGTGGATACGGGCAGGCTAGAGGCTGGTAGGGGC<br>AAGCGGAATTCCTGGTGTAGCGGTGAAATGCGCAGATATCAGGAGGAACACCGGTGGCGAAGGCGG<br>CTTGCTGGGCCGGTTCTGACGCTGAGGAGCGAGAGCGTGGGGAGCGAACAGG |
| Bacteria | Actinobacteria | Rubrobacteria | Rubrobacterales | Rubrobacteriaceae | Rubrobacter | 6 | 0 | 0 | 6 | TACGTAGGGGGCGAGCGTTGTCCGGAATTATTGGGCGTAAAGGGCGCGTAGGCGGCCCGGTAAGTCT<br>GCTGTGAAACCTTGGGGCTCAACCCCAAGCGTGCAGTGGATACTGTCGGGCTAGAGGATGGTAGAGG<br>CCAGTGGAATTCCCGGTGTAGCGGTGAAATGCGCAGATATCGGGAGGAACACCAGTAGCGAAGGCGG<br>CTGGCTGGGCCATTCCTGACGCTGAGGCGCGAAAGCTAGGGGAGCGAACAGG |
| Bacteria | Bacteria | Bacteria | Bacteria | Bacteria | Bacteria | 368 | 0 | 368 | 0 | TACGTAGGGGGCAAACGTTGTTCGGAATCACTGGGCGTAAAGGGTATGTAGGTTGTTAAGTAAGTAA<br>AAGGTGAAATCCCAAAGCTCAACTTTGGAATTGCTTTTTATACTGCTTAACTAGGGTCTCTTATAGAGGT<br>AGTAGGAATTCCTAGTGGAGGAGTAAAATCTGTAGATATTAGGAGGAACACCAATGGCGAAGGCAAG<br>CTACTGGGTAGAGACTGACATTGAGATACGAAAGCTAGGGGAGCAAATGGG |
| Bacteria | Bacteria | Bacteria | Bacteria | Bacteria | Bacteria | 98 | 0 | 98 | 0 | TACGAGTGCCCCGAGCGTTGTCCGGAATTACTGGGCGTAAAGGGTGCGTAGGGGGCGCTGTTAGTCCT<br>GTTTTAAATCCTGGTGCTCAACATCAGGAGTGGGCGGGAAACGGCAGCGCTTAGAGGACGCGAGAGG<br>TCTGCGGAACTCACGGTGTAGGGGTGAAATGCATAGATATTACACAGAACACCGATAGCGAAGGCAGC<br>TCACTAGGCCTGGATTGACGCTCAGGGACGAAAGCGTGGGGATCAAACAGG |
| Bacteria | Bacteroidetes | Bacteroidia | Bacteroidales | Bacteroidaceae | Bacteroides | 262 | 0 | 0 | 262 | TACGGAGGATCCGAGCGTTATCCGGATTTATTGGGTTTAAAGGGAGCGTAGGTGGACAGTTAAGTCA<br>GTTGTGAAAGTTTGCGGCTCAACCGTAAAATTGCAGTTGATACTGGCTGTCTTGAGTACAGTAGAGGT<br>GGGCGGAATTCGTGGTGTAGCGGTGAAATGCTTAGATATCACGAAGAACTCCGATTGCGAAGGCAGC<br>TCACTGGACTGCAACTGACACTGATGCTCGAAAGTGTGGGTATCAAACAGG |
| Bacteria | Bacteroidetes | Bacteroidia | Bacteroidales | Bacteroidaceae | Bacteroides | 251 | 0 | 0 | 251 | TACGGAGGATCCGAGCGTTATCCGGATTTATTGGGTTTAAAGGGAGCGTAGGTGGATTGTTAAGTCA<br>GTTGTGAAAGTTTGCGGCTCAACCGTAAAATTGCAGTTGATACTGGATATCTTGAGTGCAGTTGAGGC<br>AGGCGGAATTCGTGGTGTAGCGGTGAAATGCTTAGATATCACGAGGAACTCCGATTGCGAAGGCAGC<br>TCACTGGACTGCAACTGACACTGATGCTCGAAAGTGTGGGTATCAAACAGG |
| Bacteria | Bacteroidetes | Bacteroidia | Bacteroidales | Bacteroidaceae | Bacteroides | 57 | 0 | 0 | 57 | TACGGAGGACCCGAGCGTTATCCGGATTTATTGGGTTTAAAGGGAGCGTAGGTGGACAGTTAAGTCA<br>GTTGTGAAAGTTTGCGGCTCAACCGTAAAATTGCAGTTGATACTGGCTGTCTTGAGTACAGTAGAGGT<br>GGGCGGAATTCGTGGTGTAGCGGTGAAATGCTTAGATATCACGAAGAACTCCGATTGCGAAGGCAGC<br>TCACTGGACTGCAACTGACACTGATGCTCGAAAGTGTGGGTATCAAACAGG |
| Bacteria | Bacteroidetes | Bacteroidia | Bacteroidia | Bacteroidia | Bacteroidia | 115 | 0 | 115 | 0 | TACGAGTGCCCCGAGCGTTGTCCGGAATTACTGGGCGTAAAGGGTGCGTAGGGGGCGCTGTTAGTCCT<br>GTTTTAAATCCTGGTGCTCAACATCAGGAGTGGGCGGGAAACGGCAGCGCTTAGAGGACGCGAGAGG<br>TCTGCGGAACTCACGGTGTAGGGGTGAAATGCATAGATATTACACAGAACACCGATAGCGAAGGCAGC<br>TCACTAGGCCTGGATTGACGCTCAGGGACGAAAGCGTGGGGAGCAAACAGG |
| Bacteria | Bacteroidetes | Bacteroidia | Chitinophagales | Chitinophagaceae | Chitinophaga | 343 | 0 | 0 | 343 | TACGGAGGGTGCAAGCGTTATCCGGATTCATTGGGTTTAAAGGGTGCGTAGTCGGATTTGTAAGTCCG<br>TGGTGAAATCTCCAGGCTTAACCTGGAAACTGCCGTGGATACTATAAATCTTGAATGTTGTGGAGGTTA<br>GCGGAATATGTCATGTAGCGGTGAAATGCATAGATATGACATAGAACACCAATTGCGAAGGCAGCTGG<br>CTACACAAATATTGACGCTGAGGCACGAAAGCGTGGGGATCAAACAGG |
| Bacteria | Bacteroidetes | Bacteroidia | Chitinophagales | Chitinophagaceae | Chitinophaga | 333 | 0 | 0 | 333 | TACGGAGGGTGCAAGCGTTATCCGGATTCACTGGGTTTAAAGGGTGCGTAGGCGGATTTGTAAGTCC<br>GTGGTGAAATCTCCAGGCTTAACCTGGAAACTGCCGTGGATACTATAAATCTTGAATGTTGTGGAGGTT<br>AGCGGAATATGTCATGTAGCGGTGAAATGCATAGATATGACATAGAACACCAATTGCGAAGGCAGCTG<br>GCTACACAAATATTGACGCTGAGGCACGAAAGCGTGGGGATCAAACAGG |
| Bacteria | Bacteroidetes | Bacteroidia | Cytophagales | Hymenobacteraceae | Hymenobacter | 179 | 0 | 179 | 0 | TACGGAGGGTGCGAGCGTTGTCCGGATTTATTGGGTTTAAAGGGTGCGTAGGCGGCTCATTAAGTCC<br>GGGGTGAAAGCCCGTTGCTCAACAACGGAACTGCCCTGGATACTGGTGAGCTTGAGTACAGACGAGGT<br>TGGCGGAATGGACCGAGTAGCGGTGAAATGCATAGATACGGTCCAGAACCCCGATTGCGAAGGCAGC<br>TGACTAGGCTGTTACTGACGCTGAGGCACGACAGCGTGGGGAGCGAACAGG |
| Bacteria | Bacteroidetes | Bacteroidia | Cytophagales | Microscillaceae | Microscillaceae | 42 | 0 | 42 | 0 | TACGTAGGTGGCAAGCGTTGTCCGGATTTATTGGGTTTAAAGGGTGCGTAGGCGGTCTATTAAGTCAG<br>TGGTGAAATACTCTAGCTCAACTAGAGGGGTGCCATTGATACTGATGGACTTGAGTACAGATGAGGTA<br>GGCGGAATTGACGGTGTAGCGGTGAAATGCTTAGATATCGTCCAGAACACCGATAGCGAAGGCAGCTT<br>ACTAAGGAGTAACTGACGCTGAGGCACGAAAGTGTGGGGATCAAACAGG |
| Bacteria | Bacteroidetes | Bacteroidia | Flavobacteriales | Crocinitomicaceae | Fluviicola | 29 | 29 | 0 | 0 | TACGGAGGGTGCAAGCGTTGTCCGGATTTATTGGGTTTAAAGGGTGCGCAGGTGGTTTATTAAGTCA<br>GTGGTGAAAGACGGTCGCTTAACGATAGAACTGCCATTGATACTGTAAGTCTTGAATTCGGTCGGAGT<br>GGGCGGAATGTGTAGTGTAGCGGTGAAATGCATAGATATTACACAGAACACCGATAGCGAAGGCAGC<br>TCACTAGGCCTGAATTGACGCTCAGGGACGAAAGCGTGGGGATCAAACAGG |
| Bacteria | Bacteroidetes | Bacteroidia | Flavobacteriales | Flavobacteriaceae | Flavobacteriaceae | 16 | 16 | 0 | 0 | TACGGAGGGTGCAAGCGTTATCCGGAATCATTGGGTTTAAAGGGTCCGTAGGCGGGCTATTAAGTCA<br>GGGGTGAAAGTTTGTGGCTCAACCATAAAATTGCCCTTGATACTGGTAGTCTTGAGTTATTGTGAAGTG<br>GTTGGAATATGTAGTGTAGCGGTGAAATGCATAGATATTACATAGAACACCGATTGCGAAGGCAGATC<br>ACTAACAATCGACTGACGCTGAGGTGCGAAAGTGTGGGGAGCAAACAGG |
| Bacteria | Bacteroidetes | Bacteroidia | Flavobacteriales | Flavobacteriaceae | Kordia | 369 | 0 | 0 | 369 | TACGGAGGATCCAAGCGTTATCCGGAATCATTGGGTTTAAAGGGTCCGCAGGCTGTTGTTTAAGTCAG<br>AGGTGAAAGTTTGCAGCTCAACTGTAAAATTGCCTTTGATACTGGATGACTTGAGTTATAATGAAGTGG<br>TTAGAATATGTAGTGTAGCGGTGAAATGCATAGATATTACATAGAATACCGATTGCGAAGGCAGATCA<br>CTAATTATAAACTGACGCTGAGGGACGAAAGCGTGGGGAGCGAACAGG |
| Bacteria | Bacteroidetes | Bacteroidia | Flavobacteriales | Flavobacteriaceae | Kordia | 74 | 0 | 0 | 74 | TACGGAGGACCCAAGCGTTATCCGGAATCATTGGGTTTAAAGGGTCCGCAGGCTGTTGTTTAAGTCAG<br>AGGTGAAAGTTTGCAGCTCAACTGTAAAATTGCCTTTGATACTGGATGACTTGAGTTATAATGAAGTGG<br>TTAGAATATGTAGTGTAGCGGTGAAATGCATAGATATTACATAGAATACCGATTGCGAAGGCAGATCA<br>CTAATTATAAACTGACGCTGAGGGACGAAAGCGTGGGGAGCGAACAGG |

| | | | | | | | | | | |
|---|---|---|---|---|---|---|---|---|---|---|
| Bacteria | Cyanobacteria | Oxyphotobacteria | Synechococcales | Cyanobiaceae | Prochlorococcus_MIT9313 | 260 | 0 | 260 | 0 | TACGGGAGTGGCAAGCGTTATCCGGAATTATTGGGCGTAAAGCGTCCGCAGGCGGCTTTTCAAGTCTGCTGTTAAAGCGTGGAGCTTAACTCCATCATGGCAGTGGAAACTGAAAGGCTTGAGTATGGTAGGGGCAGAGGGAATTCCCGGTGTAGCGGTGAAATGCGTAGATATCGGGAAGAACACCAGTGGCGAAGGCGCTCTGCTGGGCCATTACTGACGCTCATGGACGAAAGCCAGGGGAGCGAAAGGG |
| Bacteria | Firmicutes | Bacilli | Bacillales | Bacillaceae | Bacillus | 10 | 0 | 0 | 10 | TACGTAGGTGGCAAGCGTTGTCCGGAATTATTGGGCGTAAAGCGCGCGCAGGCGGTTCCTTAAGTCTGATGTGAAAGCCCCCGGCTCAACCGGGGAGGGTCATTGGAAACTGGGGAACTTGAGTGCAGAAGAGGAGAGCGGAATTCCACGTGTAGCGGTGAAATGCGTAGAGATGTGGAGGAACACCAGTGGCGAAGGCGGCTCTCTGGTCTGTAACTGACGCTGAGGCGCGAAAGCGTGGGGAGCGAACAGG |
| Bacteria | Firmicutes | Bacilli | Bacillales | Family_X | Thermicanus | 2 | 0 | 0 | 2 | TACGTAGGGGGCGAGCGTTGTCCGGAATGATTGGGCGTAAAGCGCGCGCAGGCGGTCCTTTAAGTCTGATGTAAAAGCCCGCGGCTTAACCGCGGAAGGTCATTGGAAACTGGGGGACTTGAGGCTAGGAGAGGGAAGTGGAATTCCTGGTGTAGCGGTGAAATGCGTAGAGATCAGGAGGAATACCGATGGCGAAAGCAACTTCCTGGCTTAGAACTGACGCTGATGCGCGAAAGCGTGGGGAGCAAACAGG |
| Bacteria | Firmicutes | Bacilli | Bacillales | Paenibacillaceae | Brevibacillus | 8 | 0 | 0 | 8 | TACGTAGGTGGCAAGCGTTGTCCGGATTTATTGGGCGTAAAGCGCGCGCAGGCGGCTATGTAAGTCTGGTGTTAAAGCCCGGAGCTCAACTCCGGTTCGCATCGGAAACTGTGTAGCTTGAGTGCAGAAGAGGAAAGCGGTATTCCACGTGTAGCGGTGAAATGCGTAGAGATGTGGAGGAACACCAGTGGCGAAGGCGGCTTTCTGGTCTGTAACTGACGCTGAGGCGCGAAAGCGTGGGGAGCAAACAGG |
| Bacteria | Firmicutes | Bacilli | Lactobacillales | Streptococcaceae | Streptococcus | 55 | 0 | 0 | 55 | TACGTAGGTCCCGAGCGTTGTCCGGATTTATTGGGCGTAAAGCGAGCGCAGGCGGTTAGATAAGTCTGAAGTTAAAGGCTGTGGCTTAACCATAGTACGCTTTGGAAACTGTTTAACTTGAGTGCAAGAGGGGAGAGTGGAATTCCATGTGTAGCGGTGAAATGCGTAGATATATGGAGGAACACCGGTGGCGAAAGCGGCTCTCTGGCTTGTAACTGACGCTGAGGCTCGAAAGCGTGGGGAGCAAACAGG |
| Bacteria | Firmicutes | Clostridia | Clostridiales | Family_XII | Fusibacter | 180 | 0 | 180 | 0 | TACGTAGGGGGCAAGCGTTATCCGGAATCACTGGGCGTAAAGGGAGTGTAGGCGGTCAAGCAAGTCAGAGGTGAAAGGCATCGGCTCAACCGGTGTAAGCCTTTGAAACTGTTTGACTTGAGTGCAGGAGAGGTAAGTGGAATTCCTAGTGTAGCGGTGAAATGCGTAGATATTAGGAGGAACACCAGTGGCGAAGGCGACTTACTGGACTGTAACTGACGCTGAGGCTCGAAAGCGTGGGGAGCGAACAGG |
| Bacteria | Firmicutes | Clostridia | Clostridiales | Ruminococcaceae | Ruminococcaceae_UCG-005 | 20 | 0 | 20 | 0 | TACGTAGGTTGCAAGCGTTGTCCGGATTTACTGGGTGTAAAGGGCGTGTAGGCGGAGATGCAAGTTGGGAGTGAAATCCCGGGGCTCAACCCCGGAACTGCTCTCAAAACTGTATCCCTTGAGTATCGGAGAGGCAAGCGGAATTCCTAGTGTAGCGGTGAAATGCGTAGATATTAGGAGGAACACCAGTGGCGAAGGCGGCTTGCTGGACGACAACTGACGCTGAGGCGCGAAAGCGTGGGGAGCAAACAGG |
| Bacteria | Planctomycetes | Planctomycetacia | Pirellulales | Pirellulaceae | Pir4_lineage | 141 | 0 | 141 | 0 | TACGAACTGTGCGAACGTTATTCGGAATCACTGGGCTTAAAGGGTTTGTAGGCGGCCTTGTAAGTCAGGTGTGAAAGCCCTCGGCTCAACCGAGGAACAGCGCTTGATACTGCAAGGCTTGAGGGAGACAGGGGTAAGCGGAACTGATGGTGGAGCGGTGAAATGCGTTGATATCATCAGGAACACCGGTGGCGAAAGCGGCTTACTGGGTCTCTTCTGACGCTGAGGCGCGAAAGCCAGGGGAGCAAACGGG |
| Bacteria | Planctomycetes | Planctomycetacia | Pirellulales | Pirellulaceae | Pirellulaceae | 23 | 23 | 0 | 0 | TACAGAGGGGGCAAGCGTTGTTCGGAATTACTGGGCGTAAAGGGCGCGTAGGCGGCCCTCTAAGTCAGACGTGAAATCCCTCGGCTCAACCGAGGAACGGCGCTTGATACTGCAAGGCTTGAGGGAGACAGGGGTAAGCGGAACTGATGGTGGAGCGGTGAAATGCGTTGATATCATCAGGAACACCGGTGGCGAAAGCGGCTTACTGGGTCTCTTCTGACGCTGAGGAACGAAAGCTAGGGTAGCGAACGGG |
| Bacteria | Proteobacteria | Alphaproteobacteria | Acetobacterales | Acetobacteraceae | Rhodovarius | 16 | 0 | 0 | 16 | TACGAAGGGGGCTAGCGTTACTCGGAATGACTGGGCGTAAAGGGCGCGTAGGCTGTTTGGTAAGTTGGGCGTGAAATTCCTGGGCTTAACCTGGGGGCTGCGTCCAAGACTGCCTGACTTGAGGATGGAAGAGGCTCGTGGAATTCCCAGTGTAGAGGTGAAATTCGTAGATATTGGGAAGAACACCGGTGGCGAAGGCGGCGAGCTGGTCCATTACTGACGCTGAGGCGCGATAGCGTGGGGAGCAAACAGG |
| Bacteria | Proteobacteria | Alphaproteobacteria | Caulobacterales | Caulobacteraceae | Brevundimonas | 14 | 0 | 14 | 0 | TACGAAGGGGGCTAGCGTTGCTCGGAATTACTGGGCGTAAAGGGAGCGTAGGCGGACATTTAAGTCAGGGGTGAAATCCCAGAGCTCAACTCTGGAACTGCCTTTGATACTGGGTGTCTTGAGTGTGATAGAGGTATGTGGAACTCCGAGTGTAGAGGTGAAATTCGTAGATATTCGGAAGAACACCAGTGGCGAAGGCGACATACTGGATCATTACTGACGCTGAGGCTCGAAAGCGTGGGGAGCAAACAGG |
| Bacteria | Proteobacteria | Alphaproteobacteria | Caulobacterales | Caulobacteraceae | Caulobacter | 12 | 0 | 12 | 0 | TACGAAGGGGGCTAGCGTTGCTCGGAATTACTGGTCGTAAAGGGAGCGTAGGCGGACTGTTTAGTCAGAGGTGAAAGCCCAGGGCTCAACCTTGGAATTGCCTTTGATACTGGCAGTCTTGAGTACGGAAGAGGTATGTGGAACTCCGAGTGTAGAGGTGAAATTCGTAGATATTCGGAAGAACACCAGTGGCGAAGGCGACATACTGGTCCGTTACTGACGCTGAGGCTCGAAAGCTTGGGGAGCAAACAGG |
| Bacteria | Proteobacteria | Alphaproteobacteria | Rhizobiales | Amb-16S-1323 | Amb-16S-1323 | 76 | 0 | 0 | 76 | TACGAAGGGGGCTAGCGTTGCTCGGAATCACTGGGCGTAAAGCGCACGTAGGTGGGTCATTTAGTCAGGGGTGAAATCCTGGAGCTCAACTCCAGAACTGCCTTTGATACTGATGATCTTCGAGTCCGGGAGAGGTGGGTGGAACTGCGAGTGTAGAGGTGAAATTCGTAGATATTCGCAAGAACACCAGTGGCGAAGGCGGCTCACTGGCCCGGTACTGACGCTGAGGTGCGAAAGCGTGGGGAGCAAACAGG |
| Bacteria | Proteobacteria | Alphaproteobacteria | Rhizobiales | Devosiaceae | Devosia | 14 | 0 | 0 | 14 | TACGAAGGGGGCTAGCGTTGTTCGGATTTACTGGGCGTAAAGCGCACGTAGGCGGTTTGTTAAGTCAGGGGTGAAATCCCGGAGCTCAACTCCGGAACTGCCTTTGATACTGGCAAGCTAGAGTCCGGAAGAGGTAAGTGGAACTCCTAGTGTAGAGGTGGAATTCGTAGATATTAGGAAGAACACCAGTGGCGAAGGCGGCTTACTGGTCCGGAACTGACGCTGAGGTGCGAAAGCGTGGGGAGCAAACAGG |
| Bacteria | Proteobacteria | Alphaproteobacteria | Rhizobiales | Methyloligellaceae | Methyloligellaceae | 184 | 0 | 0 | 184 | TACGGAGGGGACTAGCGTTGTTCGGAATTACTGGGCGTAAAGCGCACGTAGGCGGATTTGTAAGTCAGGGGTGAAATCCCGAGGCTCAACCTCGGAACTGCCTTTGATACTGCAAATCTCGAGTCCGGAAGAGGTGAGTGGAATTCCTAGTGTAGAGGTGAAATTCGTAGATATTAGGAAGAACACCAGTGGCGAAGGCGGCTCACTGGTCCGGTACTGACGCTGAGGTGCGAAAGCGTGGGGAGCAAACAGG |
| Bacteria | Proteobacteria | Alphaproteobacteria | Rhizobiales | Rhizobiaceae | Candidatus_Liberibacter | 53 | 0 | 53 | 0 | TACGAAGGGGGCGAGCGTTGTTCGGAATAACTGGGCGTAAAGGGCGCGTAGGCGGGCGATTAAGTTAGAGGTGAAATCCCAGGGCTCAACCTTGGAACTGCCTTTAATACTGGTTGTCTAGAGTTTAGGAGAGGTGAGTGGAATTCCGAGTGTAGAGGTGAAATTCGTAGATATTCGGAGGAACACCGGTGGCGAAGGCGGCTCACTGGCCTGATACTGACGCTGAGGCGCGAAAGCGTGGGGAGCAAACAGG |

| | | | | | | | | | | |
|---|---|---|---|---|---|---|---|---|---|---|
| Bacteria | Proteobacteria | Alphaproteobacteria | Rhizobiales | Rhizobiales | Rhizobiales | 305 | 0 | 0 | 305 | TACGGAGGGGACTAGCGTTGTTCGGAATTACTGGGCGTAAAGCGCACGTAGGCGGATTTGTAAGTCA<br>GGGGTGAAATCCCGAGGCTCAACCTCGGAACTGCCTTTGATACTGCAAGTCTCGAGTCCGGAAGAGGT<br>GAGTGGAATTCCTAGTGTAGAGGTGAAATTCGTAGATATTAGGAAGAACACCAGTGGCGAAGGCGGC<br>TCACTGGTCCGGTACTGACGCTGAGGTGCGAAAGCGTGGGGAGCAAACAGG |
| Bacteria | Proteobacteria | Alphaproteobacteria | Rhizobiales | Rhizobiales | Rhizobiales | 225 | 0 | 0 | 225 | TACGAAGGGGGCTAGCGTTGTTCGGATTTACTGGGCGTAAAGCGCGTGTAGGCGGGTCGTTAAGTTG<br>GGGGTGAAATCCCGAGGCTCAACCTCGGAACTGCCTTTGATACTGCAAATCTCGAGTCCGATAGAGGTG<br>AGTGGAACTGCGAGTGTAGAGGTGAAATTCGTAGATATTCGCAAGAACACCAGTGGCGAAGGCGGCT<br>CACTGGTCCGGTACTGACGCTGAGGTGCGAAAGCGTGGGGAGCAAACAGG |
| Bacteria | Proteobacteria | Alphaproteobacteria | Rhizobiales | Xanthobacteraceae | Bradyrhizobium | 446 | 0 | 0 | 446 | TACGAAGGGGGCTAGCGTTGCTCGGAATCACTGGGCGTAAAGGGTGCGTAGGCGGGTCTTTAAGTCA<br>GGGGTGAAATCCTGGAGCTCAACTCCAGAACTGCCTTTGATACTGAAGATCTTGAGTTCGGGAGAGGT<br>GAGTGGAACTGCGAGTGTAGAGGTGAAATTCGTAGATATTCGCAAGAACACCAGTGGCGAAGGCGGC<br>TCACTGGCCCGATACTGACGCTGAGGTGCGAAAGCGTGGGGAGCAAACAGG |
| Bacteria | Proteobacteria | Alphaproteobacteria | Rhizobiales | Xanthobacteraceae | Xanthobacteraceae | 415 | 0 | 415 | 0 | TACGAAGGGGGCTAGCGTTGTTCGGAATCACTGGGCGTAAAGGGTGCGTAGGCGGGTCTTTAAGTCA<br>GGGGTGAAATCCTGGAGCTCAACTCCAGAACTGCCTTTGATACTGGCGATCTTGAGTTCGGGAGAGGT<br>GAGTGGAACTGCGAGTGTAGAGGTGAAATTCGTAGATATTCGCAAGAACACCAGTGGCGAAGGCGGC<br>TACCTGGCCCAATACTGACACTCAGGTGCGACAGCGTGGGGAGCAAACAGG |
| Bacteria | Proteobacteria | Alphaproteobacteria | Rhizobiales | Xanthobacteraceae | Xanthobacteraceae | 161 | 0 | 0 | 161 | TACGAAGGGTGCAAGCGTTACTCGGAATCACTGGGCGTAAAGCGCACGTAGGCGGATCGTTAAGTCA<br>GGGGTGAAATCCTGGAGCTCAACTCCAGAACTGCCTTTGATACTGGCGACCTTGAGTTCGGGAGAGGT<br>GAGTGGAACTGCGAGTGTAGAGGTGAAATTCGTAGATATTCGCAAGAACACCAGTGGCGAAGGCGGC<br>TCACTGGCCCGATACTGACGCTGAGGTGCGAAAGCGTGGGGAGCAAACAGG |
| Bacteria | Proteobacteria | Alphaproteobacteria | Rhizobiales | Xanthobacteraceae | Xanthobacteraceae | 128 | 0 | 0 | 128 | TACGAAGGGGGCTAGCGTTGCTCGGAATCACTGGGCGTAAAGCGCACGTAGGCGGATCGTTAAGTCA<br>GGGGTGAAATCCTGGAGCTCAACTCCAGAACTGCCTTTGATACTGGCGACCTTGAGTTCGGGAGAGGT<br>GAGTGGAACTGCGAGTGTAGAGGTGAAATTCGTAGATATTCGCAAGAACACCAGTGGCGAAGGCGGC<br>TCACTGGCCCGATACTGACGCTGAGGTGCGAAAGCGTGGGGAGCAAACAGG |
| Bacteria | Proteobacteria | Alphaproteobacteria | Rhizobiales | Xanthobacteraceae | Xanthobacteraceae | 41 | 0 | 0 | 41 | TACGAAGGGCGCAAGCGTTACTCGGAATCACTGGGCGTAAAGCGCACGTAGGCGGATCGTTAAGTCA<br>GGGGTGAAATCCTGGAGCTCAACTCCAGAACTGCCTTTGATACTGGCGACCTTGAGTTCGGGAGAGGT<br>GAGTGGAACTGCGAGTGTAGAGGTGAAATTCGTAGATATTCGCAAGAACACCAGTGGCGAAGGCGGC<br>TCACTGGCCCGATACTGACGCTGAGGTGCGAAAGCGTGGGGAGCAAACAGG |
| Bacteria | Proteobacteria | Alphaproteobacteria | Rhodobacterales | Rhodobacteraceae | Rhodobacteraceae | 316 | 0 | 316 | 0 | TACGGAGGGGGTTAGCGTTGTTCGGAATTACTGGGCGTAAAGCGCGCGTAGGCGGATTGGAAAGTTA<br>GAGGTGAAATCCCGGGGCTCAACCCCGGAACTGCCTTTAAAACTCCCAGTCTTGAGTTCGAGAGAGGTG<br>AGTGGAACTCCGAGTGTAGAGGTGAAATTCGTAGATATTCGGAAGAACACCAGTGGCGAAGGCGGCT<br>CACTGGCTCGATACTGACGCTGAGGTGCGAAAGTGTGGGGAGCAAACAGG |
| Bacteria | Proteobacteria | Alphaproteobacteria | Rhodobacterales | Rhodobacteraceae | Rhodobacteraceae | 300 | 0 | 0 | 300 | TACGGAGGGGGTTAGCGTTGTTCGGAATTACTGGGCGTAAAGCGCGCGTAGGCGGACCAGAAAGTTG<br>GGGGTGAAATCCCGGGGCTCAACCCCGGAACTGCCTCCAAAACTCCTGGTCTAGAGTTCGAGAGAGGT<br>GAGTGGAATTCCGAGTGTAGAGGTGAAATTCGTAGATATTCGGAGGAACACCAGTGGCGAAGGCGGC<br>TCACTGGCTCGATACTGACGCTGAGGTGCGAAAGTGTGGGGAGCAAACAGG |
| Bacteria | Proteobacteria | Alphaproteobacteria | Rhodobacterales | Rhodobacteraceae | Rhodobacteraceae | 152 | 0 | 0 | 152 | TACGGAGGGGGTTAGCGTTGTTCGGAATTACTGGGCGTAAAGCGCGCGTAGGCGGATTGGAAAGTT<br>GGGGGTGAAATCCCGGGGCTCAACCCCGGAACTGCCTCCAAAACTCCCAGTCTCGAGGATGAGAGAGG<br>CAAGTGGAATACCGAGTGTAGAGGTGAAATTCGTAGATATTCGGTGGAACACCCGTGGCGAAGGCGG<br>CTTGCTGGCTCATTTCTGACGCTGAGGTGCGAAAGCGTGGGGAGCAAACAGG |
| Bacteria | Proteobacteria | Alphaproteobacteria | Rhodobacterales | Rhodobacteraceae | Rhodobacteraceae | 141 | 0 | 0 | 141 | TACGGAGGGGGCTAGCGTTGTTCGGAATTACTGGGCGTAAAGCGCGCGTAGGCGGACTGGAAAGTT<br>GGGGGTGAAATCCCAGGGCTCAACCCTGGAACTGCCTTCAAAACTCCCAGTCTCGAGGATGAGAGAGG<br>CAAGTGGAATACCGAGTGTAGAGGTGAAATTCGTAGATATTCGGTGGAACACCCGTGGCGAAGGCGG<br>CTTGCTGGCTCATTTCTGACGCTGAGGTGCGAAAGCGTGGGGAGCAAACAGG |
| Bacteria | Proteobacteria | Alphaproteobacteria | Rickettsiales | Midichloriaceae | MD3-55 | 1242 | 32 | 1210 | 0 | TACGAAGGGGGCAAGCGTTACTCGGAATTATTGGGCGTAAAGCGTGCGTAGGCGGTTTTATAAGTTG<br>AAAGTGAAAGCCTTTGGCTCAACCAAAGAATTGCTTACAAAACTGTAAAACTAGAGTATTAGAGAAGAT<br>AGAAGAATTCCTGATGTAGGGGTGAAATCCGTAGATATCAGGAGGAATATCGAAGGCGAAAGCATCT<br>GTCTGGCTAAATACTGACGCTGTTGCACGAAAGCGTGGGGAGCAAACAGG |
| Bacteria | Proteobacteria | Alphaproteobacteria | Rickettsiales | Midichloriaceae | MD3-55 | 405 | 0 | 405 | 0 | TACGAAGGGAGCAAGCGTTACTCGGAATTATTGGGCGTAAAGCGTGCGTAGGCGGTTTTATAAGTTG<br>AAAGTGAAAGCCTTTGGCTCAACCAAAGAATTGCTTACAAAACTGTAAAACTAGAGTATTAGAGAAGAT<br>AGAAGAATTCCTGATGTAGGGGTGAAATCCGTAGATATCAGGAGGAATATCGAAGGCGAAAGCATCT<br>GTCTGGCTAAATACTGACGCTGTTGCACGAAAGCGTGGGGAGCAAACAGG |
| Bacteria | Proteobacteria | Alphaproteobacteria | Rickettsiales | Midichloriaceae | MD3-55 | 195 | 0 | 195 | 0 | TACGAAGGGGGCAAGCGTTACTCGGAATTATTGGGCGTAAAGCGTGCGTAGGCGGTTTTATAAGTTG<br>AAAGTGAAAGCCTTTGGCTCAACCAAAGAATTGCTTACAAAACTGTAAAACTAGAGTATTAGAGAGGAT<br>AGAAGAATTCCTGATGTAGGGGTGAAATCCGTAGATATCAGGAGGAATATCGAAGGCGAAAGCATCT<br>GTCTGGCTAAATACTGACGCTGTTGCACGAAAGCGTGGGGAGCAAACAGG |
| Bacteria | Proteobacteria | Alphaproteobacteria | Rickettsiales | Midichloriaceae | MD3-55 | 78 | 0 | 78 | 0 | TACGAAGGGGGCAAGCGTTACTCTGAATTATTGGGCGTAAAGCGTGCGTAGGCGGTTTTATAAGTTGA<br>AAGTGAAAGCCTTTGGCTCAACCAAAGAATTGCTTACAAAACTGTAAAACTAGAGTATTAGAGAAGATA<br>GAAGAATTCCTGATGTAGGGGTGAAATCCGTAGATATCAGGAGGAATATCGAAGGCGAAAGCATCTG<br>TCTGGCTAAATACTGACGCTGTTGCACGAAAGCGTGGGGAGCAAACAGG |
| Bacteria | Proteobacteria | Alphaproteobacteria | Sphingomonadales | Sphingomonadaceae | Altererythrobacter | 24 | 0 | 0 | 24 | TACGGAGGGAGCTAGCGTTGTTCGGAATTACTGGGCGTAAAGCGCGCGTAGGCGGCTATTCAAGTCA<br>GAGGTGAAATCCCGGGGCTCAACCCCGGAACTGCCTTTGAAACTGGATGGCTAGAATCCTGGAGAGGC<br>GAGTGGAATTCCGAGTGTAGAGGTGAAATTCGTAGATATTCGGAAGAACACCAGTGGCGAAGGCGAC<br>TCGCTGGACAGGTATTGACGCTGAGGTGCGAAAGCGTGGGGAGCAAACAGG |

| | | | | | | | | | | |
|---|---|---|---|---|---|---|---|---|---|---|
| Bacteria | Proteobacteria | Deltaproteobacteria | Myxococcales | bacteriap25 | bacteriap25 | 6 | 0 | 0 | 6 | TACAGAGGGCGCTAGCGTTGTTCGGAATCATTGGGCGTAAAGGGCGTGTAGGCGGTTTGCTAAGTCATGTGTGAAATCCCTCGGCTCAACCGGGGAACGACGCCTGAAACTGGCAAGCTAGAGTACCAAAGAGGGGGGTGGAATTCCCGGTGTAGCGGTGAAATGCGTAGATATCGGGAGGAACACCTGTGGCGAAGGCGGCCCCCTGGTTGGATACTGACGCTGAGACGCGAAAGCGTGGGGAGCAAACAGG |
| Bacteria | Proteobacteria | Deltaproteobacteria | Myxococcales | Myxococcales | Myxococcales | 339 | 0 | 0 | 339 | TACGAAGGGGGCGAGCGTTGTTCGGATTTACTGGGCGTAAAGGGCGCGCAGGCGGCGGACCAAGTCAGGTGTGAAAGCCCCAGGCTTAACCTGGGAAGTGCACTCGAAACTGCAGGGCTTGAGTATGGAAGAGGGTCTCGGAATTCCCGGTGTAGAGGTGAAATTCGTAGATATCGGGAGGAACACCAGTGGCGAAGGCGGAGACCTGGGCCAATACTGACGCTGAGGTGCGAAAGCGTGGGGAGCAAACAGG |
| Bacteria | Proteobacteria | Deltaproteobacteria | Myxococcales | P3OB-42 | P3OB-42 | 21 | 0 | 21 | 0 | TACAGAGGGCGCAAACGTTGCTCGGAATGACTGGGCGTAAAGCGCGCGTAGGCGGTGGGTTAAGTCGAATGTGAAAGCCCATGGCTCAACCATGGAAGCGCATTCGAAACTGGCTCACTGGAGTCCCGGAGAGGGTGGTGGAATTCCTAGTGTAGAGGTGAAATTCGTAGAGATTAGGAGGAACACCGGTGGCGAAGGCGACCATCTGGACGGGTACTGACGCTGAGGCGCGAAAGCGTGGGTAGCGAACAGG |
| Bacteria | Proteobacteria | Gammaproteobacteria | Alteromonadales | Shewanellaceae | Shewanella | 264 | 0 | 264 | 0 | TACGGAGGGTGCGAGCGTTAATCGGAATTACTGGGCGTAAAGCGTGCGCAGGCGGTTTGTTAAGCGAGATGTGAAAGCCCCGGGCTCAACCTGGGAACCGCATTTCGAACTGGCAAACTAGAGTCTTGTAGAGGGGGGTAGAATTCCAGGTGTAGCGGTGAAATGCGTAGAGATCTGGAGGAATACCGGTGGCGAAGGCGGCCCCCTGGACAAAGACTGACGCTCAGGCACGAAAGCGTGGGGAGCAAACAGG |
| Bacteria | Proteobacteria | Gammaproteobacteria | Alteromonadales | Shewanellaceae | Shewanella | 110 | 0 | 110 | 0 | TACGGAGGGTGCGAGCGTTAATCGGAATTACTGGGCGTAAAGCGTGCGCAGGCGGTTTGTTAAGCGAGATGTGAAAGCCCCGGGCTCAACCTGGGAACCGCATTTCGAACTGGCAAACTAGAGTATTGTAGAGGGGGGTAGAATTCCAGGTGTAGCGGTGAAATGCGTAGAGATCTGGAGGAATACCGGTGGCGAAGGCGGCCCCCTGGACAAAGACTGACGCTCAGGCACGAAAGCGTGGGGAGCAAACAGG |
| Bacteria | Proteobacteria | Gammaproteobacteria | Betaproteobacteriales | Burkholderiaceae | Acidovorax | 93 | 0 | 0 | 93 | TACGTAGGGTGCAAGCGTTAATCGGAATTACTGGGCGTAAAGCGTGCGCAGGCGGTTATATAAGACAGATGTGAAATCCCCGGGCTCAACCTGGGAACTGCATTTGTGACTGTATAGCTAGAGTACGGCAGAGGGGGATGGAATTCCGCGTGTAGCAGTGAAATGCGTAGATATGCGGAGGAACACCGATGGCGAAGGCAATCCCCTGGGCCTGTACTGACGCTCATGCACGAAAGCGTGGGGAGCAAACAGG |
| Bacteria | Proteobacteria | Gammaproteobacteria | Betaproteobacteriales | Burkholderiaceae | Burkholderia-Caballeronia-Paraburkholderia | 295 | 0 | 0 | 295 | TACGTAGGGTGCGAGCGTTAATCGGAATTACTGGGCGTAAAGCGTGCGCAGGCGGTGATGTAAGACCGATGTGAAATCCCCGGGCTTAACCTGGGAACTGCATTGGTGACTGCATCGCTGGAGTATGGCAGAGGGGGTAGAATTCCACGTGTAGCAGTGAAATGCGTAGAGATGTGGAGGAATACCGATGGCGAAGGCAGCCCCCTGGGCCAATACTGACGCTCATGCACGAAAGCGTGGGGAGCAAACAGG |
| Bacteria | Proteobacteria | Gammaproteobacteria | Betaproteobacteriales | Burkholderiaceae | Burkholderia-Caballeronia-Paraburkholderia | 245 | 0 | 0 | 245 | TACGTAGGGTGCAAGCGTTAATCGGAATTACTGGGCGTAAAGCGTGCGCAGGTGGTTATGTAAGACAGATGTGAAATCCCCGGGCTTAACCTGGGAACTGCATTGGTGACTGGCGGGCTGGAGTATGGCAGAGGGGGGTAGAATTCCACGTGTAGCAGTGAAATGCGTAGAGATGTGGAGGAATACCGATGGCGAAGGCAGCCCCCTGGGCCAATACTGACGCTCATGCACGAAAGCGTGGGGAGCAAACAGG |
| Bacteria | Proteobacteria | Gammaproteobacteria | Betaproteobacteriales | Burkholderiaceae | Burkholderiaceae | 263 | 0 | 0 | 263 | TACGTAGGGTGCAAGCGTTAATCGGAATTACTGGGCGTAAAGCGTGCGCAGGCGGCTTTGCAAGACAGATGTGAAATCCCCGGGCTTAACCTGGGAACTGCATTTGTGACTGCATGGCTGGAGTGTGGCAGAGGGGGATGGAATTCCGCGTGTAGCAGTGAAATGCGTAGATATCTGGAGGAACACCGGTGGTGAAGGCTGCGTCCTGGACCAATACTGACACTGAGGAGCGAAAGCCAGGTGAGCAAACGGG |
| Bacteria | Proteobacteria | Gammaproteobacteria | Betaproteobacteriales | Burkholderiaceae | Cupriavidus | 439 | 0 | 0 | 439 | TACGTAGGGTGCGAGCGTTAATCGGAATTACTGGGCGTAAAGCGTGCGCAGGCGGTTTTGTAAGACAGGCGTGAAATCCCCGAGCTCAACTTGGGAATTGCGCTTGTGACTGCAAGGCTAGAGTATGTCAGAGGGGGGTAGAATTCCACGTGTAGCAGTGAAATGCGTAGAGATGTGGAGGAATACCGATGGCGAAGGCAGCCCCCTGGGACGTCACTGACGCTCATGCACGAAAGCGTGGGGAGCAAACAGG |
| Bacteria | Proteobacteria | Gammaproteobacteria | Betaproteobacteriales | Burkholderiaceae | Cupriavidus | 44 | 0 | 0 | 44 | TACGTAGGGTGCGAGCGTTAATCGGAATTACTGGGCGTAAAGCGTGCGCAGGCGGTTTTGTAAGACAGGCGTGAAATCCCCGGGCTTAACCTGGGAATTGCGCTTGTGACTGCAAGGCTAGAGTGCGTCAGAGGGGGGTAGAATTCCACGTGTAGCAGTGAAATGCGTAGAGATGTGGAGGAATACCGATGGCGAAGGCAGCCCCCTGGGACGTGACTGACGCTCATGCACGAAAGCGTGGGGAGCAAACAGG |
| Bacteria | Proteobacteria | Gammaproteobacteria | Betaproteobacteriales | Burkholderiaceae | Polynucleobacter | 46 | 46 | 0 | 0 | TACGTAGGGTGCAAGCGTTAATCGGAATTACTGGGCGTAAAGCGTGCGCAGGCGGTTATGTAAGACAGGCGTGAAATCCCCGGGCTTAACCTGGGAATTGCGCTTGTGACTGCATAGCTAGAGTATGTCAGAGGGGGGTAGAATTCCACGTGTAGCAGTGAAATGCGTAGAGATGTGGAGGAATACCAATGGCGAAGGCAGCCCCCTGGGATAATACTGACGCTCATGCACGAAAGCGTGGGGAGCAAACAGG |
| Bacteria | Proteobacteria | Gammaproteobacteria | Betaproteobacteriales | Burkholderiaceae | Ralstonia | 20 | 0 | 0 | 20 | TACGTAGGGTCCAAGCGTTAATCGGAATTACTGGGCGTAAAGCGTGCGCAGGCGGTTGTGCAAGACCGATGTGAAATCCCCGGGCTTAACCTGGGAATTGCATTGGTGACTGCACGGCTAGAGTGTGTCAGAGGGAGGTAGAATTCCACGTGTAGCAGTGAAATGCGTAGAGATGTGGAGGAATACCGATGGCGAAGGCAGCCTCCTGGGATAACACTGACGCTCATGCACGAAAGCGTGGGGAGCAAACAGG |
| Bacteria | Proteobacteria | Gammaproteobacteria | Betaproteobacteriales | Burkholderiaceae | Tepidimonas | 11 | 0 | 11 | 0 | TACGTAGGGTGCGAGCGTTAATCGGAATTACTGGGCGTAAAGCGTGCGCAGGCGGTTGTGCAAGACAGGCGTGAAATCCCCGGGCTTAACCTGGGAATGGCGCTTGTGACTGCACGGCTGGAGTGCGGCAGAGGGGGATGGAATTCCGCGTGTAGCAGTGAAATGCGTAGATATGCGGAGGAACACCGATGGCGAAGGCAATCCCCTGGGCCTGCACTGACGCTCATGCACGAAAGCGTGGGGAGCAAACAGG |
| Bacteria | Proteobacteria | Gammaproteobacteria | Betaproteobacteriales | Nitrosomonadaceae | IS-44 | 253 | 0 | 0 | 253 | TACGTAGGGTGCAAGCGTTAATCGGAATTACTGGGCGTAAAGCGTGCGCAGGCGGCTTTGCAAGACAGATGTGAAATCCCCGGGCTTAACCTGGGAATTGCGCTTGAAACTAGCGGGCTTGAGTGCGGCAGAGGGGGGTGGAATTCCACGTGTAGCAGTGAAATGCGTAGAGATGTGGAGGAACACCGATGGCGAAGGCAGCCCCCTGGGTCGACACTGACGCTCATGCACGAAAGCGTGGGGAGCAAACAGG |
| Bacteria | Proteobacteria | Gammaproteobacteria | Betaproteobacteriales | Rhodocyclaceae | Denitratisoma | 52 | 0 | 0 | 52 | TACGTAGGGTGCGAGCGTTAATCGGAATTACTGGGCGTAAAGCGTGCGCAGGCGGCGCCATAAGACAGATGTGAAATCCCCGGGCTTAACCTGGGAACTGCGTTTGTGACTGTGGTGCTTGAGTACGGCAGAGGGGGGTGGAATTCCACGTGTAGCAGTGAAATGCGTAGAGATGTGGAGGAACACCGATGGCGAAGGCAGCCCCCTGGGCTAGTACTGACGCTCATGCACGAAAGCGTGGGTAGCAAACAGG |

| | | | | | | | | | | |
|---|---|---|---|---|---|---|---|---|---|---|
| Bacteria | Proteobacteria | Gammaproteobacteria | Enterobacteriales | Enterobacteriaceae | Enterobacteriaceae | 850 | 0 | 850 | 0 | TACGGAGGGTGCAAGCGTTAATCGGAATTACTGGGCGTAAAGCGCACGCAGGCGGTCTGTTAAGTCAGATGTGAAATCCCCGGGCTTAACCTGGGAACTGCATTTGAAACTGGCAGGCTTGAGTCTCGTAGAGGGGGGTAGAATTCCAGGTGTAGCGGTGAAATGCGTAGAGATCTGGAGGAATACCGGTGGCGAAGGCGGCCCCCTGGACGAAGACTGACGCTCAGGTGCGAAAGCGTGGGGAGCAAACAGG |
| Bacteria | Proteobacteria | Gammaproteobacteria | Enterobacteriales | Enterobacteriaceae | Enterobacteriaceae | 43 | 0 | 0 | 43 | TACGGAGGGTGCAAGCGTTAATCGGAATTACTGGGCGTAAAGCGCACGCAGGCGGTCTGTCAAGTCGGATGTGAAATCCCCGGGCTTAACCTGGGAACTGCATTCGAAACTGGCAGGCTAGAGTCTCGTAGAGGGGGGTAGAATTCCAGGTGTAGCGGTGAAATGCGTAGAGATCTGGAGGAATACCGGTGGCGAAGGCGGCCCCCTGGACGAAGACTGACGCTCAGGTGCGAAAGCGTGGGGAGCAAACAGG |
| Bacteria | Proteobacteria | Gammaproteobacteria | Enterobacteriales | Enterobacteriaceae | Lonsdalea | 15 | 0 | 0 | 15 | TACGGAGGGTGCAAGCGTTAATCGGAATGACTGGGCGTAAAGCGCACGCAGGCGGTTTGTTAAGTCAGATGTGAAATCCCCGAGCTTAACTTGGGAACTGCATTTGAAACTGGCAAGCTAGAGTCTTGTAGAGGGGGGTAGAATTCCAGGTGTAGCGGTGAAATGCGTAGAGATCTGGAGGAATACCGGTGGCGAAGGCGGCCCCCTGGACAAAGACTGACGCTCAGGTGCGAAAGCGTGGGGAGCAAACAGG |
| Bacteria | Proteobacteria | Gammaproteobacteria | Enterobacteriales | Enterobacteriaceae | Pantoea | 156 | 0 | 156 | 0 | TACGGAGGGCGCAAGCGTTAATCGGAATTACTGGGCGTAAAGCGCACGCAGGCGGTCTGTTAAGTCAGATGTGAAATCCCCGGGCTTAACCTGGGAACTGCATTTGAAACTGGCAGGCTTGAGTCTCGTAGAGGGGGGTAGAATTCCAGGTGTAGCGGTGAAATGCGTAGAGATCTGGAGGAATACCGGTGGCGAAGGCGGCCCCCTGGACGAAGACTGACGCTCAGGTGCGAAAGCGTGGGGAGCAAACAGG |
| Bacteria | Proteobacteria | Gammaproteobacteria | Gammaproteobacteria | Gammaproteobacteria | Gammaproteobacteria | 194 | 0 | 0 | 194 | TACAGAGGGTGCAAGCGTTAATCGGAATTACTGGGCGTAAAGCGTGCGCAGACGGTTGCGTAAGTCAGATGTGAAAGCCCCGGGCTCAACCTGGGAATTGCATTTGAGACTGCGTAGCTAGGGTGCGGAAGAGGGAAGCGGAATTTCCGGTGTAGCGGTGAAATGCGTAGATATCTAGAGGAACGCCTGAGGCGAAAGCGGGGCACTGGGCCGATTCTGACGCTGAGACGCGAAAGCGTGGGGAGCAAACAGG |
| Bacteria | Proteobacteria | Gammaproteobacteria | Gammaproteobacteria_Incertae_Sedis | Unknown_Family | Acidibacter | 170 | 0 | 170 | 0 | TACAGAGGGTGCGAGCGTTAATCGGAATTACTGGGCGTAAAGCGTGCGTAGACGGTTATGTAAGTCGGGTGTGAAAGCCCCGGGCTCAACCTGGGAATTGCATTCGAGACTGCATAGCTAGGGTGCGGAAGAGGGAAGCGGAATTTCCGGTGTAGCGGTGAAATGCGTAGATATCGGAAGGAACATCAGTGGCGAAAGCGGCTTCCTGGTCCAGCACCGACATTCAGGCACGAAAGCGTGGGGAGCAAACAGG |
| Bacteria | Proteobacteria | Gammaproteobacteria | Halothiobacillales | Halothiobacillaceae | Thiovirga | 21 | 21 | 0 | 0 | TACGTAGGGTGCAAGCGTTAATCGGAATTACTGGGCGTAAAGCGTGCGCAGGCGGTTATGTAAGACAGGCGTGAAATCCCCGGGCTTAACCTGGGAATTGCGCTTGTGACTGCATAGCTAGAGTGTGCAAGAGGATGGGGGAATTCCTGGTGTAGCAGTGAAATGCGTAGAGATCAGGAGGAACATCAATGGCGAAGGCACCTTTCTGGGGCAACACTGACGCTGAGAGACGAAAGCGTGGGGAGCAAACAGG |
| Bacteria | Proteobacteria | Gammaproteobacteria | Oceanospirillales | Halomonadaceae | Halomonas | 19 | 0 | 0 | 19 | TACGGAGGGTGCGAGCGTTAATCGGAATTACTGGGCGTAAAGCGCGCGTAGGCGGCGTGATAAGCCGGTTGTGAAAGCCCCGGGCTCAACCTGGGAACGGCATCCGGAACTGTCAGGCTAGAGTGCAGGAGAGGAAGGTAGAATTCCCGGTGTAGCGGTGAAATGCGTAGAGATCGGGAGGAATACCAGTGGCGAAGGCGGCCTTCTGGACTGACACTGACGCTGAGGTGCGAAAGCGTGGGTAGCAAACAGG |
| Bacteria | Proteobacteria | Gammaproteobacteria | Pseudomonadales | Moraxellaceae | Acinetobacter | 79 | 0 | 0 | 79 | TACAGAGGGTGCAAGCGTTAATCGGATTTACTGGGCGTAAAGCGCGCGTAGGTGGCTAAATAAGTCAAATGTGAAATCCCCGAGCTTAACTTGGGAATTGCATTCGATACTGTTTGGCTAGAGTATGGGAGAGGATGGTAGAATTCCAGGTGTAGCGGTGAAATGCGTAGAGATCTGGAGGAATACCGATGGCGAAGGCAGCCATCTGGCCTAATACTGACACTGAGGTGCGAAAGCATGGGGAGCAAACAGG |
| Bacteria | Proteobacteria | Gammaproteobacteria | SAR86_clade | SAR86_clade | SAR86_clade | 226 | 0 | 226 | 0 | TACGGAAGGTGCAAGCGTTAATCGGAATTACTGGGCGTAAAGCGCGCGTAGGTGGTTTGTTAAGTTGGATGTGAAAGCCCTGGGCTCAACCTAGGAACTGCATCCAAAACTAACTCACTAGAGTACGATAGAGGGAGGTAGAATTCATAGTGTAGCGGTGGAATGCGTAGATATTATGAAGAATACCAGTGGCGAAGGCGGCCTCCTGGATCTGTACTGACACTAAGGTGCGAAAGCGTGGGTAGCGAACAGG |
| Bacteria | Proteobacteria | Gammaproteobacteria | SAR86_clade | SAR86_clade | SAR86_clade | 46 | 0 | 46 | 0 | TACGGAAGGCGCAAGCGTTAATCGGAATTACTGGGCGTAAAGCGCGCGTAGGTGGTTTGTTAAGTTGGATGTGAAAGCCCTGGGCTCAACCTAGGAACTGCATCCAAAACTAACTCACTAGAGTACGATAGAGGGAGGTAGAATTCATAGTGTAGCGGTGGAATGCGTAGATATTATGAAGAATACCAGTGGCGAAGGCGGCCTCCTGGATCTGTACTGACACTAAGGTGCGAAAGCGTGGGTAGCGAACAGG |
| Bacteria | Proteobacteria | Gammaproteobacteria | Steroidobacterales | Steroidobacteraceae | Povalibacter | 260 | 0 | 0 | 260 | TACGTAGGGTGCGAGCGTTAATCGGAATTACTGGGCGTAAAGTGTGCGCAGGCGGCGCTGCAAGTCGAGTGTGAAATCCCCAGGCTTAACCTGGGAACTGCATTCGAGACTGCATTGCTAGAGTATGGGAGAGGGAAGTGGAATTTCCGGTGTAGCGGTGAAATGCGTAGATATCGGAAGGAACATCAGTGGCGAAAGCGACTTCCTGGACCAATACTGACGCTCATGTGCGAAAGCGTGGGGAGCAAACAGG |
| Bacteria | Proteobacteria | Gammaproteobacteria | Steroidobacterales | Steroidobacteraceae | Steroidobacter | 240 | 0 | 0 | 240 | TACAGAGGGTGCGAGCGTTAATCGGAATTACTGGGCGTAAAGCGCGCGTAGGCGGCTTTGCAAGTCGGGTGTGAAATCCCCAGGCTTAACCTGGGAACTGCATTCGAGACTGCATTGCTAGAGTATGGGAGAGGGAAGTGGAATTTCCGGTGTAGCGGTGAAATGCGTAGATATCGGAAGGAACATCAGTGGCGAAAGCGACTTCCTGGACCAATACTGACGCTCATGTGCGAAAGCGTGGGGAGCAAACAGG |
| Bacteria | Proteobacteria | Gammaproteobacteria | Xanthomonadales | Xanthomonadaceae | Pseudoxanthomonas | 28 | 0 | 0 | 28 | TACGAAGGGTGCAAGCGTTACTCGGAATTACTGGGCGTAAAGCGTGCGTAGGTGGTGGCTTAAGTCCGTTGTGAAAGCCCTGGGCTCAACCTGGGAATTGCAGTGGATACTGGGTCACTAGAGTGTGGTAGAGGGTGGCGGAATTCCCGGTGTAGCAGTGAAATGCGTAGAGATCGGGAGGAACACCCGTGGCGAAGGCGGCCACCTGGGCCAACACTGACACTGAGGCACGAAAGCGTGGGGAGCAAACAGG |
| Bacteria | Proteobacteria | Proteobacteria | Proteobacteria | Proteobacteria | Proteobacteria | 11 | 0 | 11 | 0 | AACTTAAGGGCTAGTTTTGTGGGCAAAAGCGTAAAGGGTGTGATAGGCCGTTTTAATTAAAAAGGGTTTTATAATTTAAGAAGGTAAATGGAATTTTTTTGTAGCGGTGGAATGTGTAAATAGAAAAAAGAACTTTAGACGTGAAGACTATTTATTTTTGAGATTATAGTGTGATGGCTAAAACACGAGAGTATGGGGAGCAAACAGG |

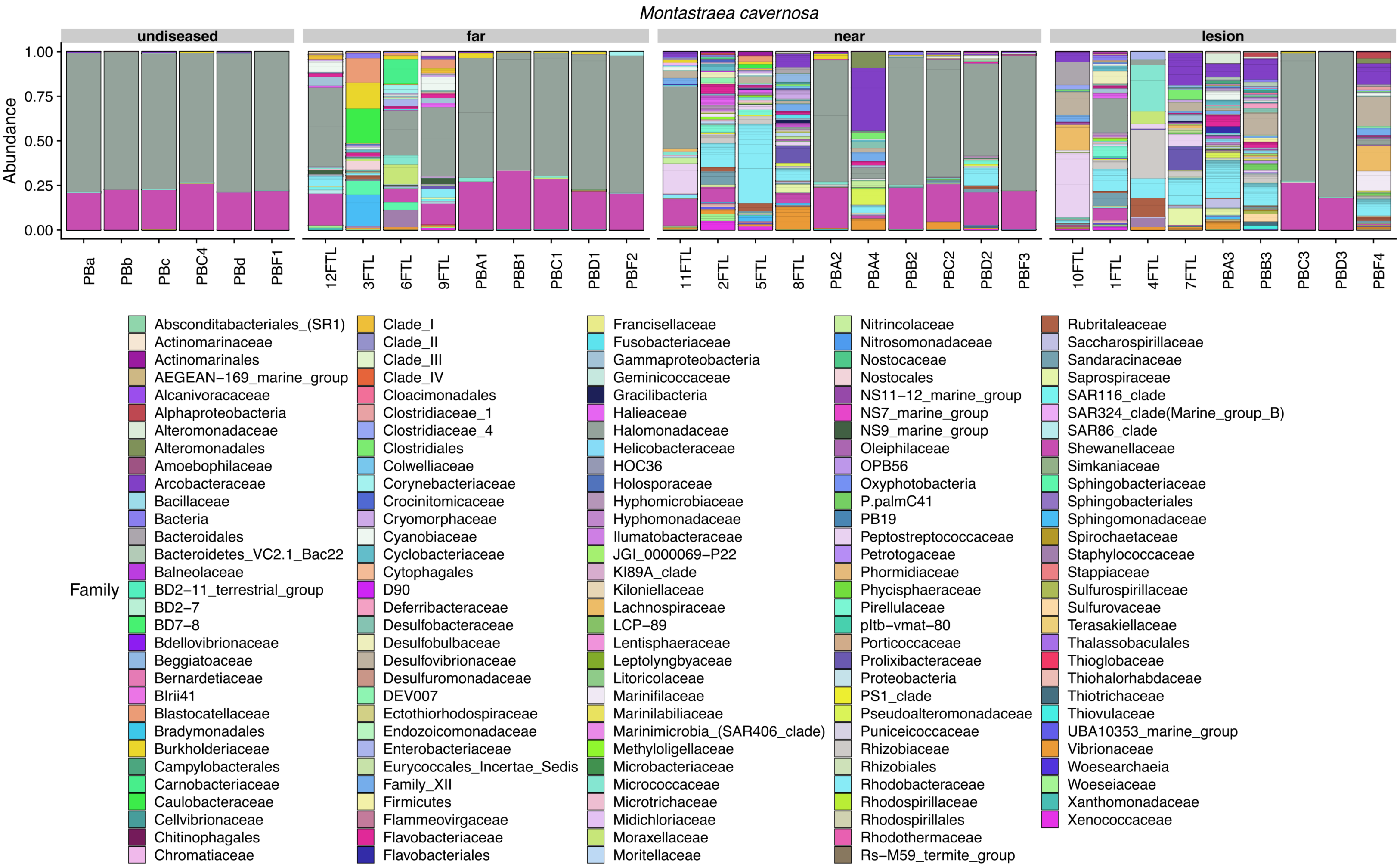


Figure S1. Relative abundance of amplicon sequence variants, colored by Family, in undiseased neighboring corals, apparently healthy tissue far from the disease lesion, apparently healthy tissue near the disease lesion, and disease lesions in *Montastraea cavernosa* with stony coral tissue loss disease. Samples with names beginning with PB were collected in July 2017 and samples with names beginning with a number were collected December 2017.

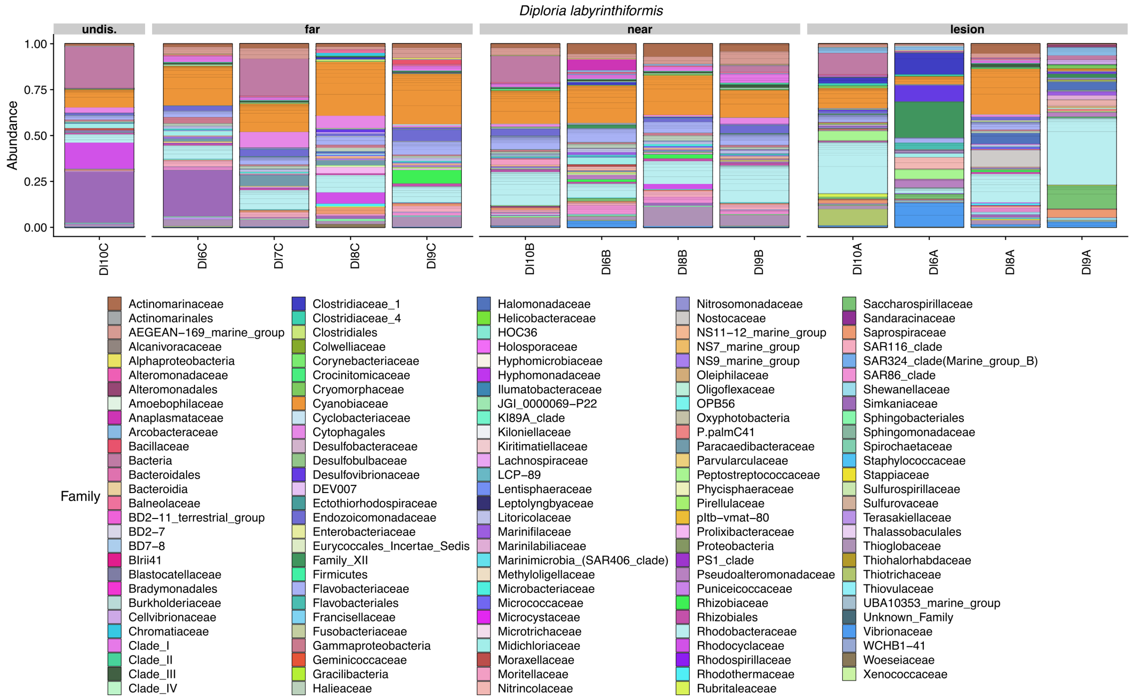


Figure S2. Relative abundance of amplicon sequence variants, colored by Family, in undiseased neighboring corals, apparently healthy tissue far from the disease lesion, apparently healthy tissue near the disease lesion, and disease lesions in *Diploria labyrinthiformis* with stony coral tissue loss disease.

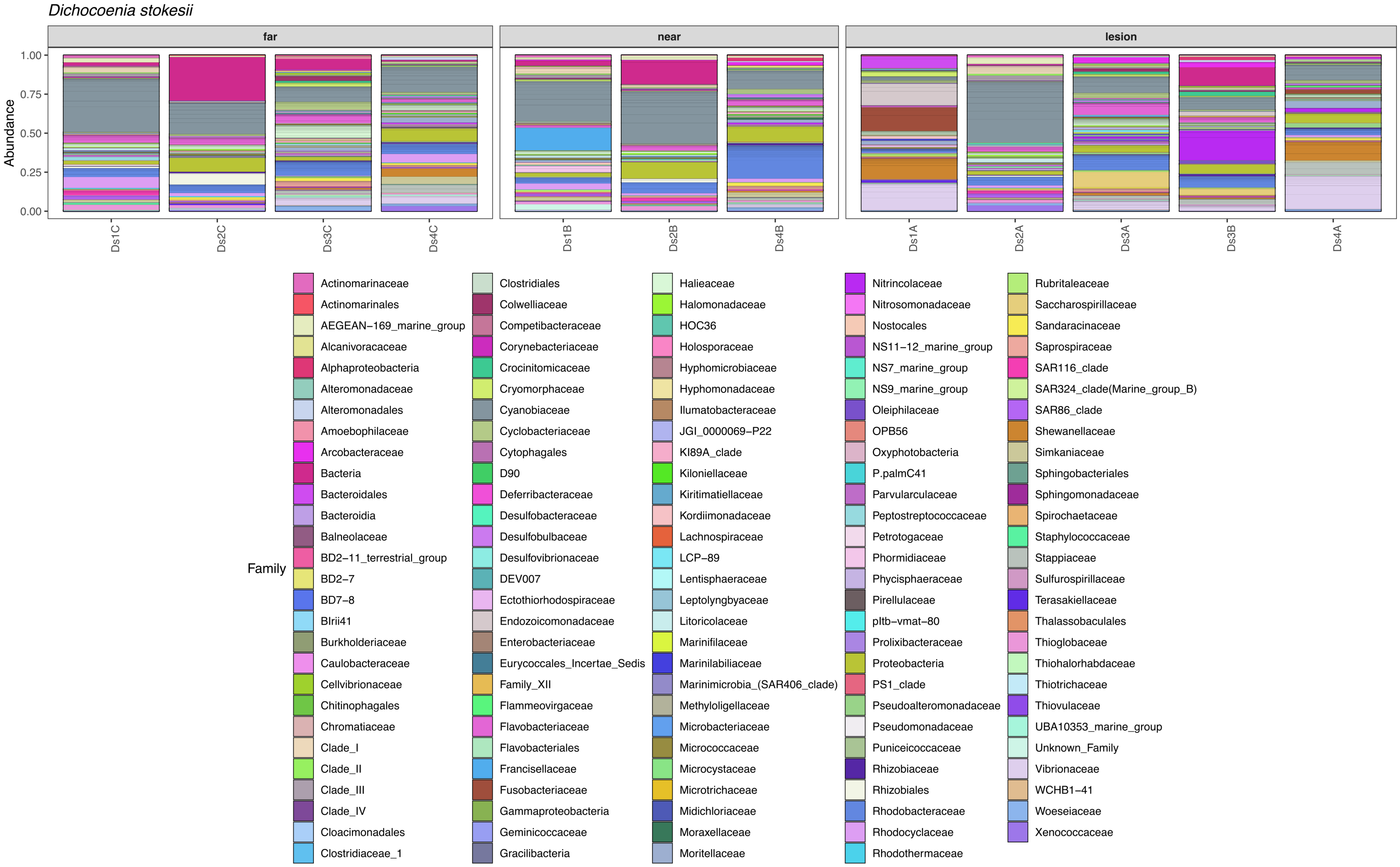


Figure S3. Relative abundance of amplicon sequence variants, colored by Family, in apparently healthy tissue far from the disease lesion, apparently healthy tissue near the disease lesion, and disease lesions in *Dichocoenia stokesii* with stony coral tissue loss disease.

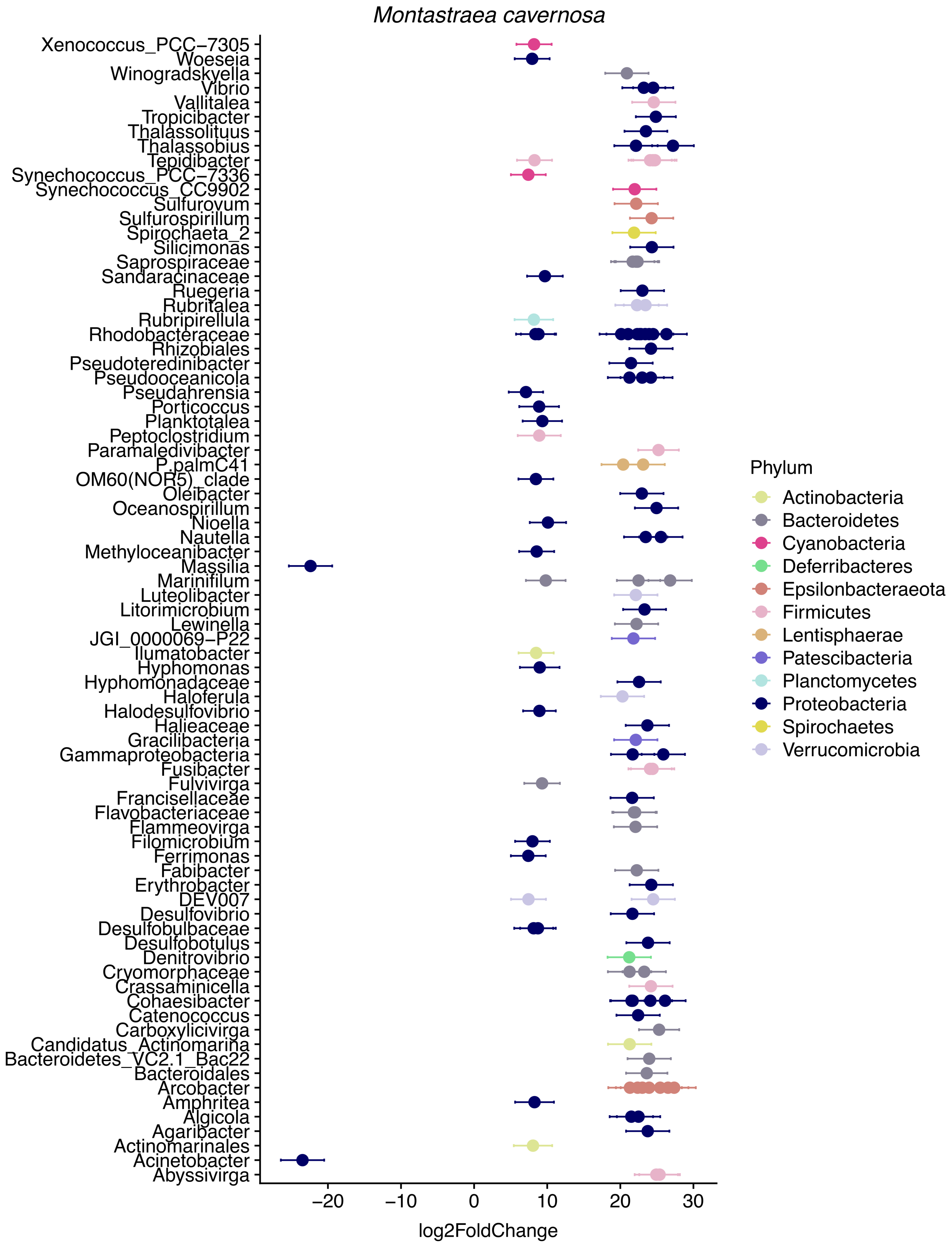


Figure S4. Differentially abundant amplicon sequence variants (ASVs) in stony coral tissue loss disease. Positive log2FoldChange values correspond to ASVs that are more abundant in disease lesions compared to apparently healthy tissue on diseased corals in *Montastraea cavernosa*.

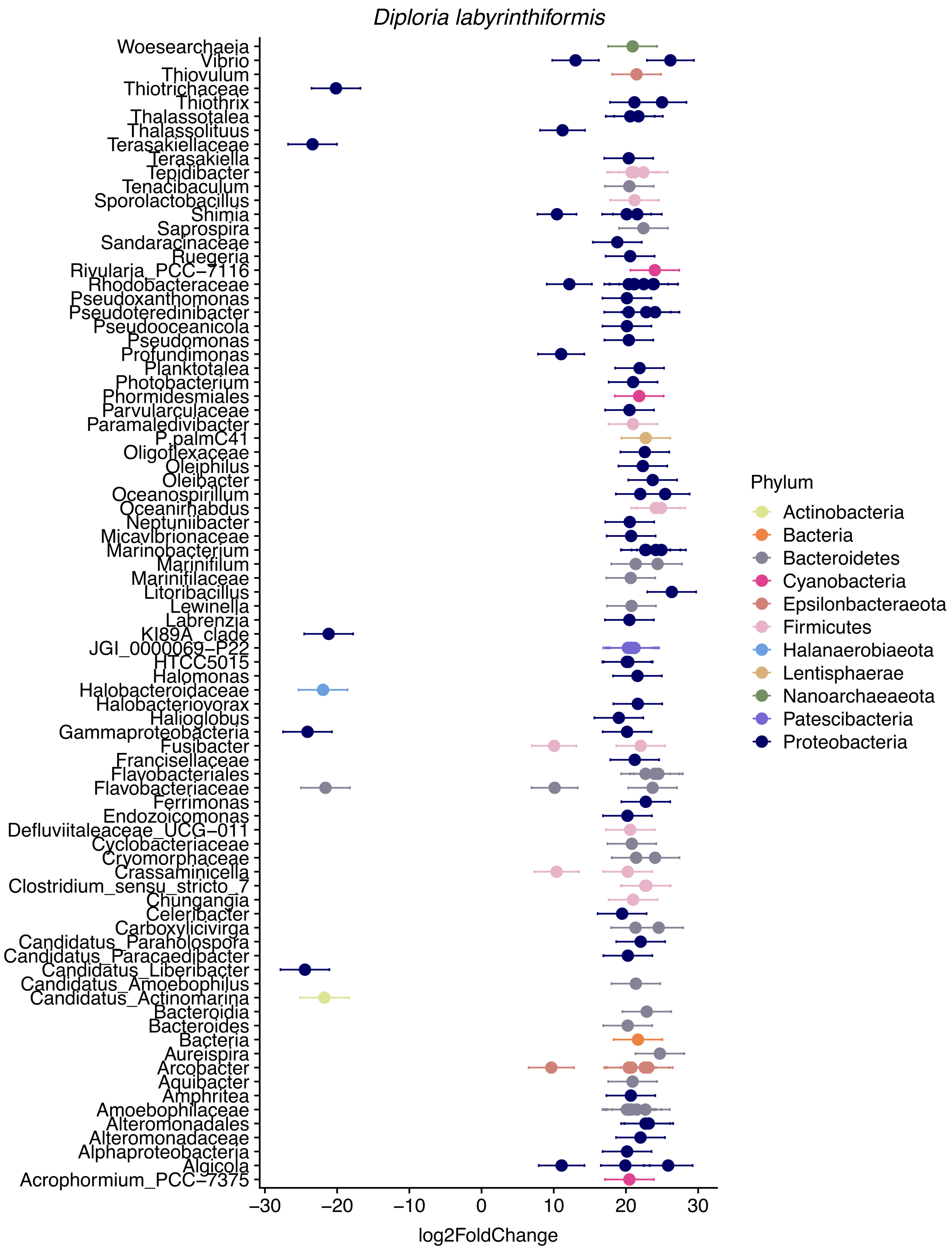


Figure S5. Differentially abundant amplicon sequence variants (ASVs) in stony coral tissue loss disease. Positive log2FoldChange values correspond to ASVs that are more abundant in disease lesions compared to apparently healthy tissue on diseased corals in *Diploria labyrinthiformis*.

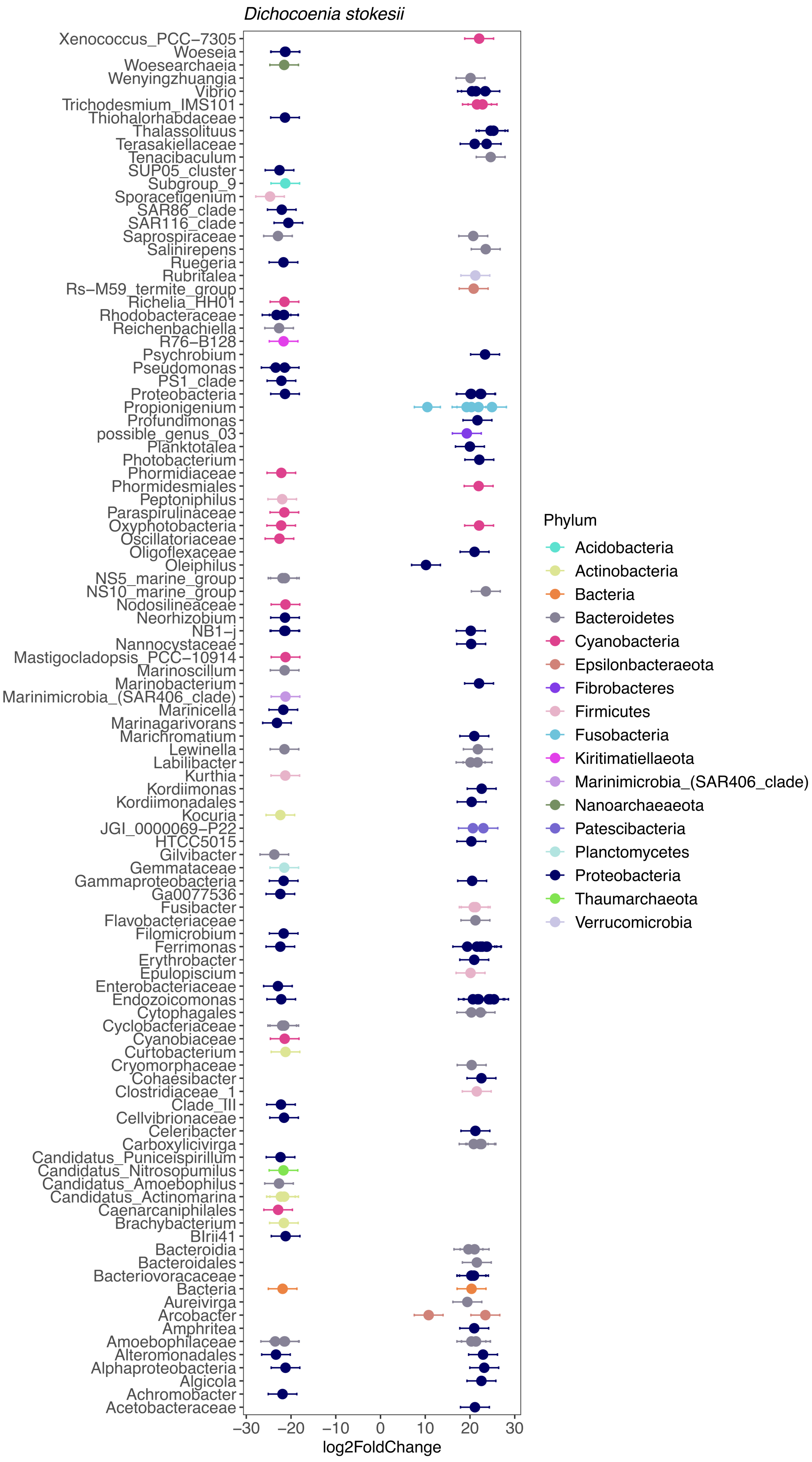


Figure S6. Differentially abundant amplicon sequence variants (ASVs) in stony coral tissue loss disease. Positive log2FoldChange values correspond to ASVs that are more abundant in disease lesions compared to apparently healthy tissue on diseased corals in *Dichocoenia stokesii*.
